# Oocyte spindle assembly depends on multiple interactions between HP1 and the CPC

**DOI:** 10.1101/2020.06.03.132142

**Authors:** Lin-Ing Wang, Tyler DeFosse, Janet K. Jang, Rachel A. Battaglia, Victoria F. Wagner, Kim S. McKim

**Affiliations:** Waksman Institute and Department of Genetics, Rutgers, the State University of New Jersey, Piscataway N.J. 08854-8020

**Keywords:** meiosis, chromosome segregation, kinetochore, oocyte, chromosome passenger complex, central spindle

## Abstract

The chromosomes in the oocytes of many animals appear to promote bipolar spindle assembly. In *Drosophila* oocytes, spindle assembly requires the chromosome passenger complex (CPC), which consists of INCENP, Borealin, Survivin and Aurora B. To determine what recruits the CPC to the chromosomes and its role in spindle assembly, we developed a strategy to manipulate the function and localization of INCENP, which is critical for recruiting the Aurora B kinase. We found that an interaction between Borealin and the chromatin is crucial for the recruitment of the CPC to the chromosomes and is sufficient to build kinetochores and recruit spindle microtubules. We also found that HP1 moves from the chromosomes to the spindle microtubules along with the CPC. We propose that the interaction with HP1 promotes the movement of the CPC from the chromosomes to the microtubules. In addition, within the central spindle, rather than at the centromeres, the CPC and HP1 are required for homologous chromosome bi-orientation.

## Introduction

Accurate chromosome segregation during cell division requires bi-orientation of homologous chromosomes in meiosis I and sister chromatids in mitosis or meiosis II. Bi-orientation is the result of two simultaneous processes: the assembly of microtubules into a bipolar spindle and the correct attachment of the kinetochores to microtubules. In mitosis and male meiosis, the bipolarity of the spindle is defined by centrosomes at each pole. These serve as microtubule organizing centers, nucleating microtubules that grow towards the chromosomes and make contact with kinetochores (Cheeseman, 2014; Heald and Khodjakov, 2015; Nicklas, 1997; Watanabe, 2012). In the oocytes of many species, the female meiotic spindle assembles without centrosomes. Spindly assembly initiates when microtubules cluster around the chromosomes after nuclear envelope breakdown (Dumont and Desai, 2012; Radford et al., 2017). In mouse oocytes, this involves the accumulation of acentriolar MTOCs around the chromosomes (Schuh and Ellenberg, 2007). In contrast, the chromosomes or chromatin serve as the sites of microtubule nucleation in the oocytes of *Drosophila* (Matthies et al., 1996; Theurkauf and Hawley, 1992), *Xenopus* (Heald et al., 1996) and human (Holubcová et al., 2015).

Chromatin-coated beads in *Xenopus* extracts (Heald et al., 1996; Sampath et al., 2004) and chromosomes without kinetochores in *Drosophila* oocytes (Radford et al., 2015) build spindles. Similarly, kinetochore-independent chromosome interactions between the chromosomes and the spindle in *C. elegans* oocyte meiosis have been observed (Dumont et al., 2010; Muscat et al., 2015; Wignall and Villeneuve, 2009). These results suggest that oocyte chromatin carries signals that can recruit and organize spindle assembly factors. Potential targets of these signals include the Ran pathway and the Chromosomal Passenger Complex (CPC), both of which have been shown to promote chromosome-directed spindle assembly (Bennabi et al., 2016; Drutovic et al., 2020; Mullen et al., 2019; Radford et al., 2017). The CPC comprises Aurora B kinase, the scaffold subunit INCENP, and the two targeting subunits Borealin and Survivin (Deterin in *Drosophila*). In *Drosophila*, the depletion of Aurora B or INCENP causes a complete failure of meiotic spindle assembly in oocytes (Colombié et al., 2008; Radford et al., 2012). Similarly, the CPC is required for promoting spindle assembly when sperm nuclei are added to *Xenopus* egg extracts (Kelly et al., 2007; Maresca et al., 2009; Sampath et al., 2004). The results in *Xenopus* and *Drosophila* suggest that oocyte chromosomes carry signals that can recruit and activate the activity of the CPC. Indeed, the *Xenopus* studies demonstrate that spindle assembly requires that the CPC interacts with both chromatin and microtubules (Tseng et al., 2010; Wheelock et al., 2017).

The CPC displays a dynamic localization pattern during cell division that contributes to its known functions. During mitosis, the CPC localizes to the centromeres during metaphase, where it is required for correcting kinetochore-microtubule (KT-MT) attachments, cohesion regulation and checkpoint regulation (Carmena et al., 2012a; Krenn and Musacchio, 2015; Trivedi and Stukenberg, 2020). It then relocates onto the microtubules to form the spindle midzone required at anaphase for cytokinesis (Adams et al., 2001; Carmena et al., 2012b; Cesario et al., 2006; Chang et al., 2006). In *Drosophila* prometaphase I oocytes, however, the CPC is most abundant on the central spindle, similar to the anaphase midzone of mitotic cells, and is not usually observed at the centromeres (Jang et al., 2005; Radford et al., 2012). Another critical component of the prometaphase meiotic spindle is Subito, a *Drosophila* Kinesin-6 and orthologue of MKLP2 (Giunta et al., 2002). Subito is required for organizing the meiotic central spindle and homologous chromosome bi-orientation in *Drosophila* oocytes (Jang et al., 2005). Subito and INCENP genetically interact and are mutually dependent for their localization (Das et al., 2018; Das et al., 2016; Radford et al., 2012). Based on these observations, a model for spindle assembly in *Drosophila* oocytes is that the CPC activates multiple spindle organizing proteins, including Subito. Hence, the goal in this study was to determine how the chromosomes are involved in this process.

To test the hypothesis that the chromosomes recruit and activate the CPC to spatially restrict oocyte spindle assembly, we developed an RNAi-resistant expression system to generate separation-of-function mutants of the CPC. The most thoroughly studied pathways for localization of the CPC to chromosomes involves two histone kinases, Haspin and Bub1, which phosphorylate H3T3 and H2AT120 (Hindriksen et al., 2017) and recruit Survivin and Borealin, respectively, to the inner centromeres. However, we provide evidence that Haspin and Bub1 are not required for spindle assembly in oocyte meiosis. Instead, an interaction between Borealin and the chromatin recruits the CPC to the oocyte chromosomes to initiate kinetochore and spindle assembly. Furthermore, Heterochromatin Protein 1 (HP1) may interact with the CPC in multiple phases of spindle assembly. HP1 colocalizes with the CPC on the chromosomes and then they both move onto the spindle. Thus, our research has revealed a mechanism for how the meiotic chromosomes recruit the CPC for spindle assembly and how the CPC moves to the microtubules. We also propose that within the central spindle, the CPC and HP1 promote the bi-orientation of homologous chromosomes in oocytes.

## Results

### Using RNAi resistant transgenes to study factors that regulate CPC localization

The *Drosophila* meiotic spindle is composed of two types of microtubules. The kinetochore microtubules (K-fibers) are defined by those that end at a kinetochore, and the central spindle is defined by microtubules that make antiparallel overlaps in the center of the spindle and contain the Kinesin 6 Subito (Jang et al., 2005). The CPC is required for the initiation of spindle assembly by recruiting kinetochore proteins to the centromeres and recruiting microtubules around the chromosomes (Radford et al., 2015; Radford et al., 2012). While the CPC localizes to the centromeres in mitotic cells prior to anaphase (Adams et al., 2001; Cesario et al., 2006; Giet and Glover, 2001), it localizes predominantly to the central spindle in prometaphase I *Drosophila* oocytes (Figure 1A). Centromere CPC localization has only been observed under certain conditions, including live imaging (Costa and Ohkura, 2019) and colchicine-treated oocytes where microtubules are destabilized (Figure 1A). Thus, the CPC can localize to meiotic chromatin in addition to spindle microtubules in metaphase oocytes.

**Fig 1:**
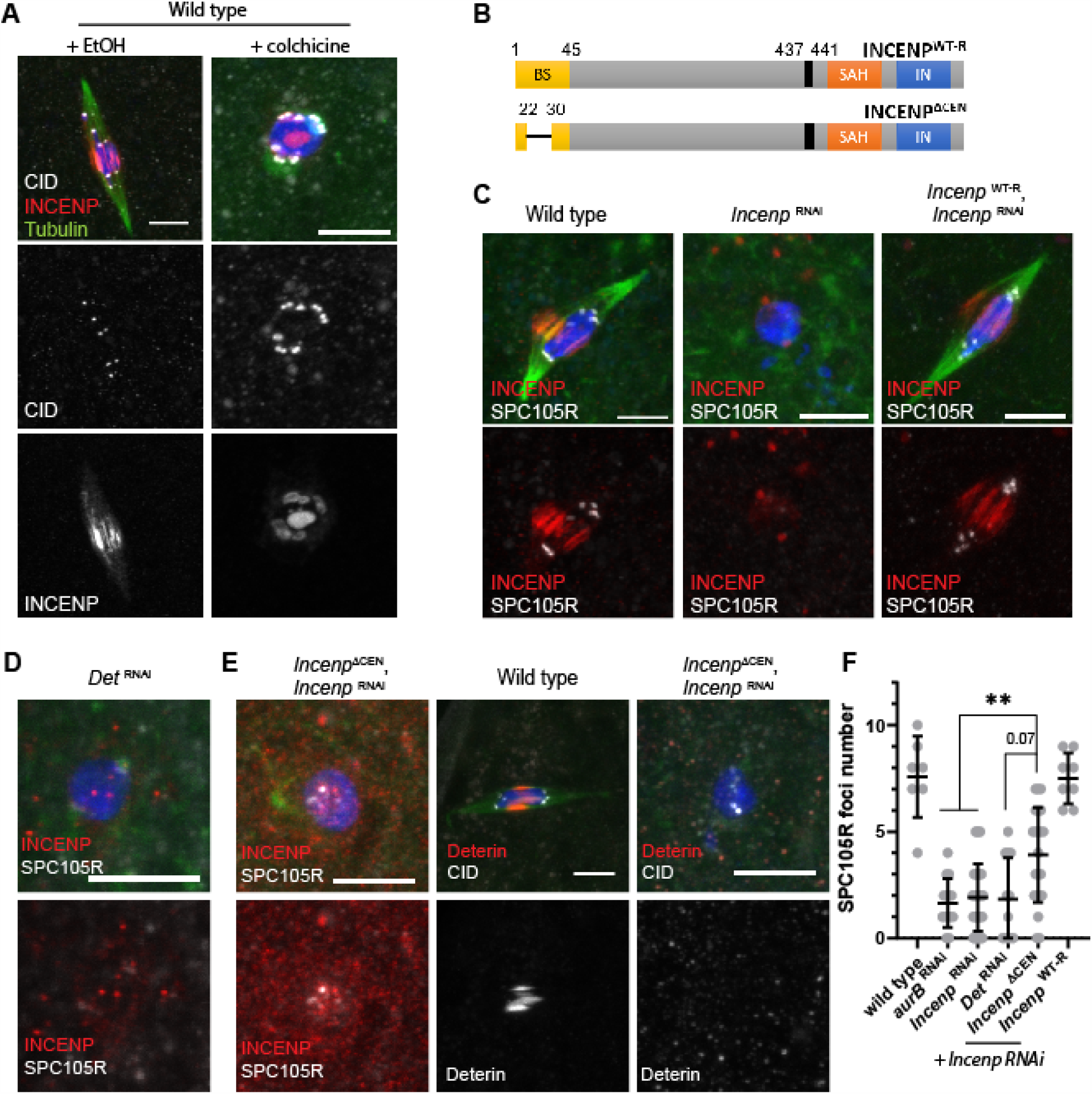
INCENP localization and spindle assembly depends on the N-terminal Borealin/Deterin binding domain. (A) INCENP localization in oocytes after a 60 minute colchicine treatment to destabilize the microtubules. The sister centromeres appeared to be separating in colchicine treated oocytes, the mechanism of which is not known. INCENP is in red, CID/CENP-A is in white, tubulin is in green and DNA is in blue. Scale bars in all images are 5 µm. (B) A schematic of *Drosophila* INCENP, showing the location of the centromere-targeting N-terminal region (BS), single alpha helix (SAH) and INbox (IN) domains. The black box (437-441) shows the sequence targeted by the shRNA *GL00279* that were mutated to make RNAi resistant *Incenp*^*WT-R*^. The conserved amino acids 22-30 are deleted in *Incenp*^*ΔCEN*^. (C) INCENP (red) localization in *Incenp* RNAi oocytes and oocytes also expressing the *Incenp*^*WT-R*^ RNAi-resistant transgene. Kinetochore protein SPC105R is in white. (D) Spindle and kinetochore assembly defects in *Deterin* RNAi oocytes. (E) Spindle and kinetochore assembly defects in *Incenp*^*ΔCEN*^, *Incenp* RNAi oocytes, with Deterin or INCENP in red and SPC105R or CID in white. (F) Quantitation of SPC105R localization in RNAi oocytes (n=7, 14, 32, 12 22 and 8 oocytes). Error bars indicate 95% confidence intervals and ** = *p*-value < 0.001 run by Fisher’s exact test.

To study the relationship between CPC localization patterns and function, we developed a system to target the CPC to distinct chromosomal or spindle locations. Oocyte-specific RNAi was used to knock down *Incenp* instead of using mutants because the CPC is essential for viability. *Incenp* or *aurB* RNAi oocytes fail to recruit kinetochore proteins, such as SPC105R or NDC80 to the centromeres, or recruit microtubules around the chromosomes (Radford et al., 2015; Radford et al., 2012). We constructed an *Incenp* transgene to be RNAi resistant with silent mismatches in the region targeted by shRNA *GL00279* (Figure 1B, Figure S 1). Expressing the RNAi-resistant transgene (*Incenp*^*WT-R*^) rescued the defects in *Incenp* RNAi oocytes, including spindle and kinetochore assembly, homolog bi-orientation, and fertility, restoring them to wild-type levels (Figure 1C, Table 1). Having successfully established this RNAi-resistant system, the *Incenp*^*WT-R*^ backbone was used to construct separation-of-function *Incenp* mutants. These mutants were analyzed in either a wild-type (i.e. *Incenp*^*WT-R*^ oocytes) or an RNAi background (i.e. *Incenp*^*WT-R*^ *Incenp RNAi* oocytes).

**Table 1:**
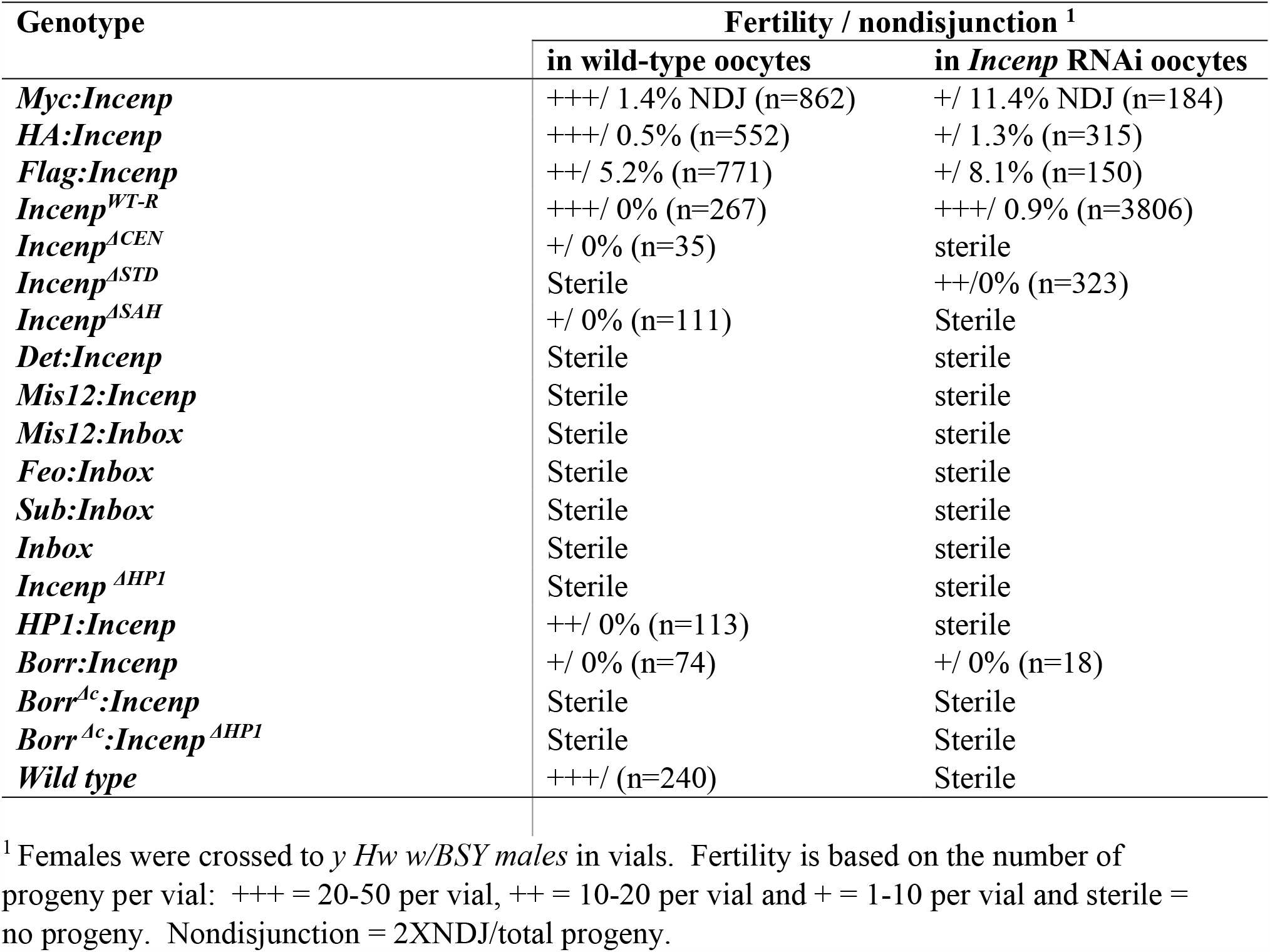
Summary of transgene fertility

### Targeting the CPC to centromeres is sufficient, but not required, to promote kinetochore microtubule (K-fiber) assembly

Borealin and Survivin (known as Deterin in *Drosophila*) targets the CPC to the centromeres in mitotic cells (Carmena et al., 2012b; Hindriksen et al., 2017). Borealin and Survivin interact with the N-terminal domain of INCENP, which is required for centromere localization of the CPC (Jeyaprakash et al., 2007; Klein et al., 2006). We were successful at generating an effective *Deterin* shRNA for oocyte RNAi and found the same phenotype as *Incenp* or *aurB* RNAi oocytes (Figure 1C and D). This result suggests that Deterin plays an important role in targeting INCENP and Aurora B to the chromosomes in *Drosophila* oocytes. To test the function of the CPC at the centromere, we deleted conserved amino acids 22-30 in INCENP (referred to as *Incenp*^*ΔCEN*^, Figure 1B, Figure S 1) that corresponds to the centromere targeting domain described in chicken INCENP and is predicted to be required for the interaction with Borealin and Deterin (Ainsztein et al., 1998; Jeyaprakash et al., 2007). In *Incenp*^*ΔCEN*^, *Incenp* RNAi oocytes, the INCENP^ΔCEN^ protein had weak localization to the chromosomes, did not recruit Deterin, or promote spindle assembly (Figure 1E). However, the *Incenp*^*ΔCEN*^, *Incenp* RNAi oocytes had an intermediate level of SPC105R localization compared to wild-type or *Incenp, aurB* RNAi or *Deterin* RNAi oocytes (Figure 1C-F). These results suggest that the N-terminal domain of INCENP recruits Deterin and is required for spindle assembly in *Drosophila* oocytes, although some kinetochore assembly is possible without it.

To directly test whether centromeric CPC can promote spindle assembly and regulate homolog bi-orientation in oocytes, we targeted the CPC to the centromeric regions. Based on a strategy used in HeLa cells, INCENP was fused to the kinetochore protein MIS12 (Liu et al., 2009). MIS12 loads onto centromeres during prophase (Schittenhelm et al., 2007; Venkei et al., 2012), is independent of other kinetochore proteins (Feijão et al., 2013; Przewloka et al., 2007) and localizes to foci on the chromosomes in wild-type and *Incenp* RNAi oocytes (Figure S 2A) (Głuszek et al., 2015). To target the CPC to the centromeres, the N-terminal amino acids 1-46 of INCENP (the “BS” domain) were replaced with MIS12 (*mis12:Incenp*) (Figure 2A).

**Fig 2:**
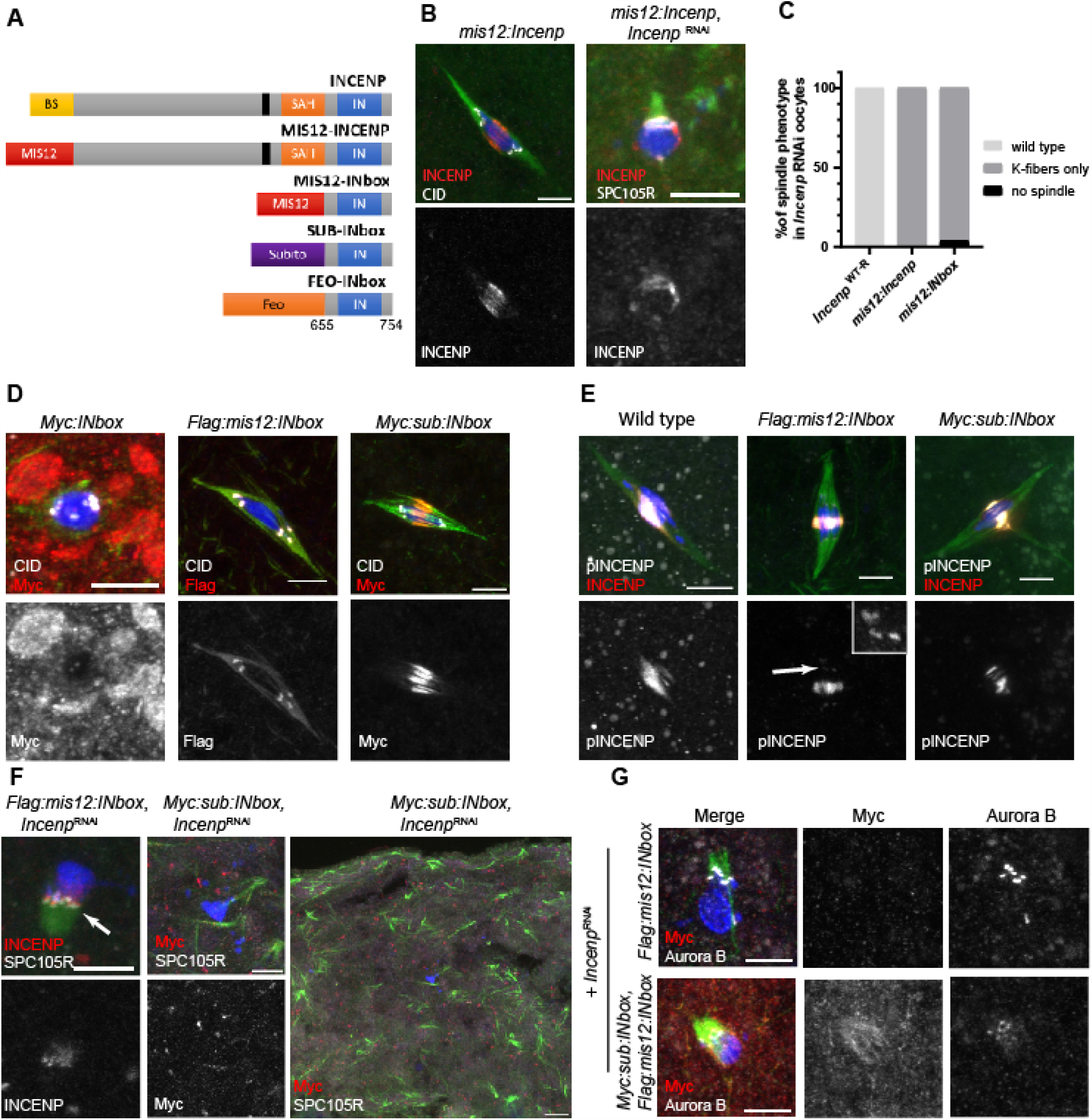
Independent localization of the CPC to the centromere and the central spindle assembles only kinetochore-dependent microtubules. (A) A schematic of the CPC constructs designed to target the CPC to the centromeres. (B) Expression of *mis12:Incenp* in wild-type and *Incenp* RNAi oocytes. (C) Quantitation of the spindle phenotype of MIS12 fusions expressed in *Incenp* RNAi oocytes (n=23, 36 and 25 oocytes). (D) *Myc:INbox, Flag:mis12:INbox* and *Myc:sub:INbox* expressed in wild-type oocytes. (E) Detection of the Aurora B substrate, phosphorylated INCENP (pINCENP). The arrow and inset shows the pINCENP signal at the centromeres when the INbox is targeted by MIS12. INCENP is in red and pINCENP is in white. (F) *Flag:mis12:INbox* and *Myc:sub:INbox* expressed in *Incenp* RNAi oocytes. The arrow points to K-fibers in *Flag:mis12:Inbox* oocytes. The low magnification image of *Myc:sub:INbox* shows microtubule bundles in the cytoplasm instead of around the chromosomes. Transgene proteins are in red, SPC105R in white, DNA is in blue and tubulin is in green. Scale bars represent 5 µm in the left three panels and 10 µm in the low magnification image. (G) *Flag:mis12:INbox* expressed alone or co-expressed with *Myc:sub:INbox* in *Incenp* RNAi oocytes. Merged images showed DNA (blue), tubulin (green), Myc (red) and Aurora B (white). Scale bars indicate 5 µm.

Surprisingly, when *mis12:Incenp* was expressed in wild-type oocytes, we did not observe centromere localization (Figure 2B). However, the females were sterile due to the failure to complete the two meiotic divisions and initiate the mitotic divisions (Figure S 2B, Table 1). This phenotype demonstrated that the transgene was expressed and toxic to the embryo. When expressing *mis12:Incenp* in *Incenp* RNAi oocytes, however, the fusion protein localized in and around the centromeres and spindle assembly was limited. Kinetochore assembly similar to wild-type and K-fiber formation was observed, which was defined as oocytes with microtubules emanating from the kinetochores (Figure 2B and C, Figure S 2C). These spindles were usually short and lacked a central spindle. These results suggest that targeting INCENP to the centromere regions could only promote kinetochore assembly and K-fiber formation.

In the presence of endogenous INCENP, MIS12:INCENP did not localize to the centromeres. To test the possibility that regions outside the BS domain of INCENP negatively regulate centromere localization, we fused MIS12 directly to the INbox domain of INCENP (amino acids 655-755 of the C-terminal domain), which is sufficient to recruit Aurora B (Figure 2A) (Bishop and Schumacher, 2002). Expressing unfused *INbox* in the presence of endogenous INCENP had a dominant negative effect on oocyte spindle assembly, causing a diminished spindle (Figure 2D) and sterility (Table 1). This observation suggests that unlocalized INbox has a dominant-negative effect. Similar to observations in mammalian cells (Gohard et al., 2014), INbox could be acting like a competitive inhibitor of INCENP by generating non-productive binding interactions with Aurora B. When expressing *mis12:INbox* in wild-type oocytes, MIS12:INbox was present at the centromeres (Figure 2D). MIS12:INbox was also observed on the spindle, although the mechanism and consequences of this are not known. Phospho-INCENP, which is a marker of Aurora B activity (Salimian et al., 2011), was observed at the central spindle and in the vicinity of the centromeres (Figure 2E), showing MIS12:INbox can successfully recruit and activate Aurora B. Similar to *mis12:Incenp*, when *mis12:INbox* was expressed in *Incenp* RNAi oocytes, SPC105R was recruited, K-fibers formed, but there was not central spindle (Figure 2C, F and G, Figure S 2C). The centromeres were unable to bi-orient and were often clustered together and oriented towards the same pole of a monopolar spindle. These results demonstrate that centromere-targeted CPC is sufficient to build kinetochores and K-fibers, consistent with findings in *Xenopus* extracts (Bonner et al., 2019), but not the central spindle.

### Independent targeting of the CPC to both the centromere and central spindle is not sufficient to assemble a wild-type spindle

Because centromere-directed Aurora B only promotes K-fiber assembly, it is possible that oocyte spindle assembly depends on microtubule-associated Aurora B. Indeed, our prior studies have suggested the CPC is simultaneously required for kinetochore and central spindle microtubule assembly in oocytes (Radford et al., 2015). Therefore, we performed experiments to determine whether the recruitment of Aurora B to these two sites is independent or whether one site might depend on the other. To target Aurora B to the spindle, the *INbox* was fused with two microtubule-associated proteins, Fascetto (*feo*, the *Drosophila* PRC1 homolog) and Subito (Figure 2A). These two fusions, *feo:INbox* and *sub:INbox*, resulted in robust INbox localization to the central spindle when expressed in wild-type oocytes (Figure 2D, E and Figure S 2D). When expressed in *Incenp* RNAi oocytes, neither *feo:INbox* nor *sub:INbox* oocytes assembled a spindle around the chromosomes (Figure 2F and Figure S 2C). These results suggest that microtubule associated Aurora B is not sufficient to promote spindle assembly around the chromosomes. Fusing Subito to the INbox promoted microtubule bundles in the cytoplasm but not in the specifically important location around the chromosomes. Additionally, most SPC105R localization was absent in *sub:INbox, Incenp* RNAi oocytes (Figure 2F, Figure S 2C) similar to *Incenp* RNAi (Figure 1C,F). One possible explanation for these observations is that the central spindle targeting of Aurora B lacked the interaction with the chromosomes necessary for spindle and kinetochore assembly.

The problem with the *feo:INbox* and *sub:INbox* experiments could have been the absence of chromosome-associated Aurora B to recruit microtubules and nucleate central spindle assembly. Therefore, to test whether independent targeting of Aurora B to the chromosomes and the microtubules would promote spindle assembly, we co-expressed *mis12:INbox* and either *sub:INbox* or *feo:INbox* in *Incenp* RNAi oocytes. Interestingly, only the K-fibers formed in these oocytes, suggesting that *sub:INbox* and *feo:INbox* cannot contribute to spindle assembly even in the presence of K-fibers (Figure 2G and Figure S 2C). We did observe enhanced microtubule bundling involving the K-fibers, indicating that SUB:INbox was active and could contribute to microtubule bundling. These results indicate that independently targeting two populations of Aurora B is not sufficient to assemble a bipolar spindle (see also Tseng et al., 2010).

### Borealin, but not Deterin, is sufficient for most meiotic spindle assembly

Central spindle assembly may require an interaction between the CPC and the chromosomes before the bundling of antiparallel microtubules. Furthermore, the phenotype of *Incenp*^*ΔCEN*^ and *Deterin* RNAi oocytes suggests that Borealin and Deterin are critical for chromosome-directed spindle assembly in oocytes (Figure 1D and E). To test whether an interaction of Deterin and/or Borealin with INCENP is sufficient to target the CPC for oocyte spindle assembly, we replaced the BS domain of INCENP with Deterin or Borealin (referred to as *Det:Incenp* and *borr:Incenp*, Figure 3A). In *Det:Incenp* oocytes, INCENP localized to the chromatin close to the centromeres and K-fibers were formed. However, DET:INCENP failed to localize to the spindle and no central spindle was observed (Figure 3B). Borealin localization could not be detected in *Det:Incenp, Incenp* RNAi oocytes (Figure S 3A), suggesting this spindle phenotype is independent of Borealin.

**Fig 3:**
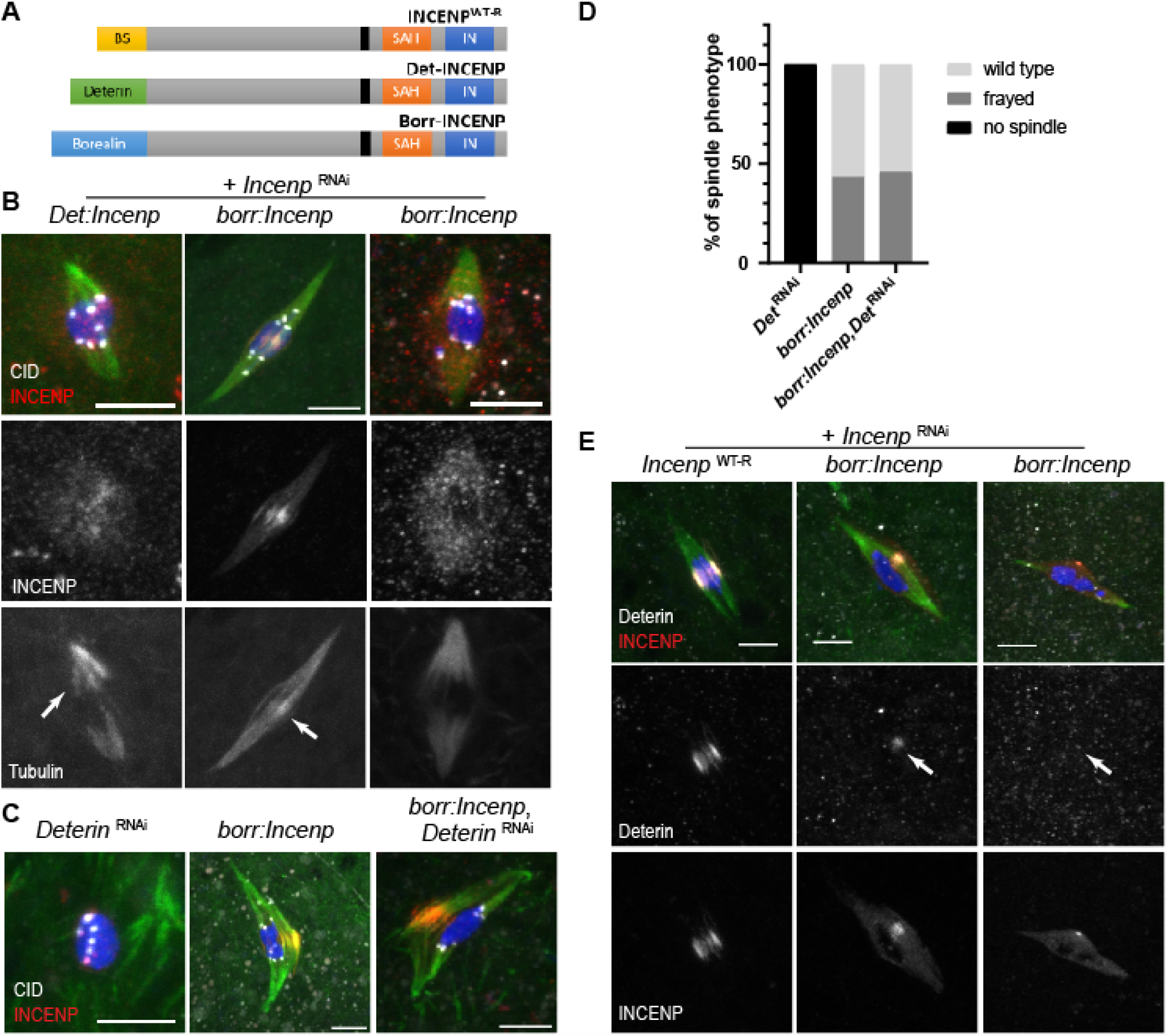
Borealin is sufficient to recruit the CPC for meiotic spindles assembling. (A) A schematic of the *Det:Incenp* and *borr:Incenp* fusions compared to wild-type *Incenp*. (B) The spindle phenotypes of *borr:Incenp* and *Det:Incenp* in *Incenp* RNAi oocytes. The separate channels show the localization of INCENP and the microtubules. The second *borr:Incenp* image is an example of a spindle with diffuse spindle INCENP. Arrows show K-fibers in *Det:Incenp* oocytes and central spindle fibers in *borr:Incenp* oocytes. INCENP is in red, DNA is in blue, tubulin is in green and CID is in white. (C, D) The effect of Deterin depletion in *borr:Incenp* oocytes (n=18, 23 and 24). (E) Deterin localization in wild-type and *borr:Incenp, Incenp* RNAi oocytes. Second *borr:Incenp* image is an example of a spindle with diffuse spindle INCENP. INCENP is in red, Deterin is in white, DNA is in blue and tubulin is in green. Scale bars in all images are 5 µm.

In contrast, *borr:Incenp* rescued spindle assembly, including the central spindle, in *Incenp* RNAi oocytes (Figure 3B) although the degree of rescue in some oocytes was variable. Some of these oocytes had frayed central spindles (Figure 3C, D) and in 47% oocytes, INCENP localized to the microtubules but failed to be concentrated in the central spindle (n=28, Figure 3B). A bipolar spindle also formed when *borr:Incenp* was expressed in *Deterin* RNAi oocytes although similar to *borr:Incenp, Incenp* RNAi oocytes, there were some spindle abnormalities (17% diffuse INCENP localization, 48% frayed spindles, Figure 3C and 3D). In contrast, 44% of *Deterin* RNAi oocytes did not assemble a spindle, and the rest only showed non-specific microtubule clustering around the chromosomes (Figure 3C and 3D). Thus, the *borr:Incenp* fusion promotes spindle assembly independent of Deterin. Deterin localized to the spindle in *borr:Incenp, Incenp* RNAi oocytes, but only when BORR:INCENP was concentrated in the central spindle (Figure 3E). These observations suggest that Borealin is sufficient to move the CPC from the chromosomes to the microtubules and promote spindle assembly in *Drosophila* oocytes. An important role for Borealin in CPC-dependent spindle assembly has also been shown in *Xenopus* (Kelly et al., 2007). Deterin, in contrast, can promote INCENP localization to the chromatin and K-fiber formation, and may have a role stabilizing the interaction of Borealin and INCENP with microtubules. Deterin is not, however, sufficient to promote movement of the CPC to the microtubules.

### Recruitment of the CPC to the chromosomes and spindle assembly depends on the C-terminal domain of Borealin

Deterin and Borealin are known to be recruited by the histone markers H3T3ph and H2AT120ph, respectively (Wang et al., 2010; Yamagishi et al., 2010). These two histones are phosphorylated by Haspin and BUB1 kinases, respectively. However, spindle assembly and CPC localization were normal in *haspin*, or *Bub1* RNAi oocytes or *haspin, Bub1* double RNAi oocytes, suggesting Haspin and BUB1 are not required for spindle assembly (Figure S 4A). In addition, fertility and chromosome segregation was not affected in any of the genotypes (Figure S 4B), ubiquitous expression of *haspin*, or *Bub1* shRNAs did not cause lethality, and *haspin* null mutants are viable and fertile (Fresan et al., 2020). These results suggest that Haspin and BUB1 are not required for the CPC to promote oocyte spindle assembly. Therefore, we investigated other mechanisms for Borealin-mediated CPC recruitment to the chromosomes.

In addition to recruitment by BUB1 activity, Borealin can be recruited to chromosomes by an interaction between its C-terminal domain and HP1 (Liu et al., 2014) or nucleosomes (Abad et al., 2019). Although the C-terminal domain of Borealin is poorly conserved, there is evidence in support of the hypothesis that the CPC interacts with HP1 during chromosome-directed spindle assembly in oocytes. INCENP and Borealin colocalize with HP1 and H3K9me3, the histone marker that recruits HP1, in *aurB* RNAi oocytes (Figure 4A and 4B). Furthermore, HP1 is present on chromosomes in *Incenp* RNAi oocytes (Figure 4A), showing that the CPC is not required for HP1 localization. To test the hypothesis that HP1 recruits the CPC to chromatin in oocytes, we deleted the C-terminal domain of Borealin from *borr:Incenp* (referred as *borr*^*ΔC*^*:Incenp*) (Figure 4C). Spindle assembly was severely impaired in *borr*^*ΔC*^*:Incenp, Incenp* RNAi oocytes. Only 19% of oocytes assembled K-fibers, and none of them assembled the central spindle (Figure 4D, 4E and 4F). Most of the oocytes that assembled K-fibers (75%) had normal SPC105R localization (Figure 4F, Figure S 3), suggesting that K-fiber formation was associated with SPC105R localization. Because CPC components and HP1 colocalize when Aurora B activity is absent, we favor the interpretation that an interaction between Borealin and HP1 is required to build both K-fibers and the central spindle in oocytes. However, we cannot rule out a role for an interaction between Borealin and the nucleosomes in recruiting the CPC to the chromosomes.

**Fig 4:**
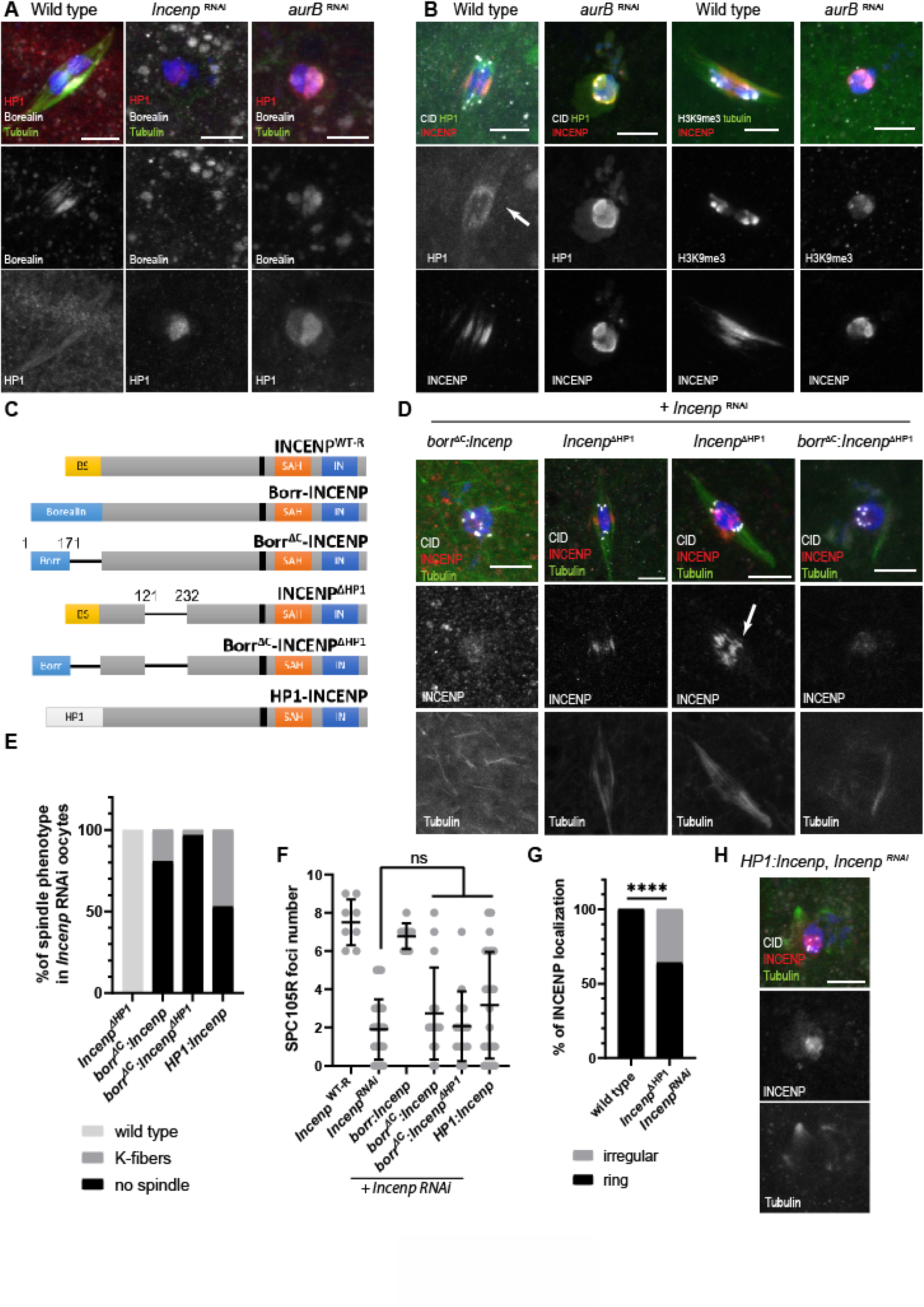
HP1-Borealin interaction is critical for CPC-dependent meiotic spindle assembly. (A) Borealin and HP1 localization in wild type, *Incenp* RNAi and *aurB* RNAi oocytes. Borealin is in white, HP1 is in red, tubulin is in green and DNA is in blue. (B) HP1 and H3K9me3 localization in wild-type and *aurB* RNAi oocytes. Arrow shows HP1 localization in wild-type oocytes on the spindle. HP1 or tubulin is in green, INCENP is in red, DNA is in blue and CID or H3K9me3 are in white. (C) A schematic of *Incenp* mutant constructs affecting HP1 interactions. Proposed HP1 binding sites are located in the C-terminus of Borealin and in INCENP between amino acids 121-232. Full length *Drosophila* HP1 was fused to INCENP by substitution for the BS domain. (D) Expression of *Incenp* transgenes shown in (C) in *Incenp* RNAi oocytes, including *borr*^*ΔC*^*:Incenp, Incenp*^*ΔHP1*^ and *borr*^*ΔC*^*:Incenp*^*ΔHP1*^. Two different images of *Incenp*^*ΔHP1*^ are shown to compare ring-shaped and disorganized (see arrow) localization of the CPC. The images show CID in white, INCENP in red, DNA in blue and tubulin in green. (E) Quantitation of spindle phenotype in the HP1-interaction-defective mutants shown in (D) (n= 17, 26, 31 and 49 oocytes, in the order as shown in the graph). (F) SPC105R localization (see Figure S 3) in the HP1-interaction-defective mutant oocytes (n= 8, 32, 9, 15, 14, 33). Bars indicates 95% confidence intervals and ** = *p-*value < 0.01 in Fisher’s exact test. (G) Quantitation of INCENP spindle localization in wild-type (n=19) and *Incenp*^*ΔHP1*^, *Incenp* RNAi oocytes (n=17). **** = *p*-value < 0.0001. (H) Expressing *HP1:Incenp* (see C) in *Incenp* RNAi oocytes. The images show CID in white, INCENP in red, DNA in blue and tubulin in green.

Two putative HP1 interaction sites exist in INCENP (Ainsztein et al., 1998; van der Horst and Lens, 2014) and these were deleted to make *Incenp*^*ΔHP1*^. While spindle assembly in *borr*^*ΔC*^*:Incenp, Incenp* RNAi was severely affected, spindle assembly in *Incenp*^*ΔHP1*^, *Incenp* RNAi oocytes was similar to wild type, except that 36% oocytes INCENP displayed irregular and disorganized central spindle localization (Figure 4D, E, and G). Thus, an INCENP-HP1 interaction may only have a minor role in oocyte spindle assembly. To test for additive effects, a mutant with all HP1 sites deleted was generated (*borr*^*ΔC*^*:Incenp*^*ΔHP1*^). A more severe spindle assembly defect was observed in *borr*^*ΔC*^*:Incenp*^*ΔHP1*^, *Incenp* RNAi oocytes. Specifically, the spindle was abolished in nearly all the oocytes and we measured a small decrease in K-fiber formation (*p* = 0.08, Figure 4D and E). These results suggest that the C-terminal domain of Borealin recruits the CPC to the oocyte chromosomes, with a minor contribution from INCENP. To test whether the only function of Borealin in oocytes is to interact with HP1 for recruitment of the CPC, the BS domain of *Incenp* was replaced with HP1 (*HP1:Incenp*) (Figure 4C). HP1:INCENP localized to part of the chromatin, probably the heterochromatin regions, but 53% of the oocytes failed at spindle assembly, and the rest only had K-fiber formation associated with SPC105R localization (Figure 4E, F and H, Figure S 3). Thus, targeting the CPC to the heterochromatin regions without Borealin is not sufficient for bipolar spindle assembly. Rather than Borealin being an adapter for CPC localization, an interaction between Borealin and HP1 and/or nucleosomes appears to be essential for transfer of the CPC to the microtubules and oocyte spindle assembly.

### Ejection of HP1 and the CPC from the chromosomes depends on Aurora B and microtubules

To determine if Borealin is sufficient to target the CPC to the chromatin, we examined the behavior of BORR:INCENP fusion proteins when Aurora B activity was inhibited. Similar to the results in *aurB* RNAi oocytes, when wild-type oocytes were treated with the Aurora B inhibitor Binucleine 2 (BN2) (Smurnyy et al., 2010), the spindle was drastically diminished and INCENP colocalized with HP1 and H3K9me3 on the chromosomes (Figure 5A). The same result was observed with *borr:Incenp, Incenp* RNAi oocytes. The BORR:INCENP fusion colocalized with HP1 on the chromosomes in BN2 treated oocytes (Figure 5A). In contrast, in *borr*^*ΔC*^*:Incenp, Incenp* RNAi oocytes, the BORR^ΔC^: INCENP fusion did not localize to the chromosomes. These results suggest the Borealin C-terminal domain is required to target the CPC to the chromosomes, including sites enriched with HP1.

**Fig 5:**
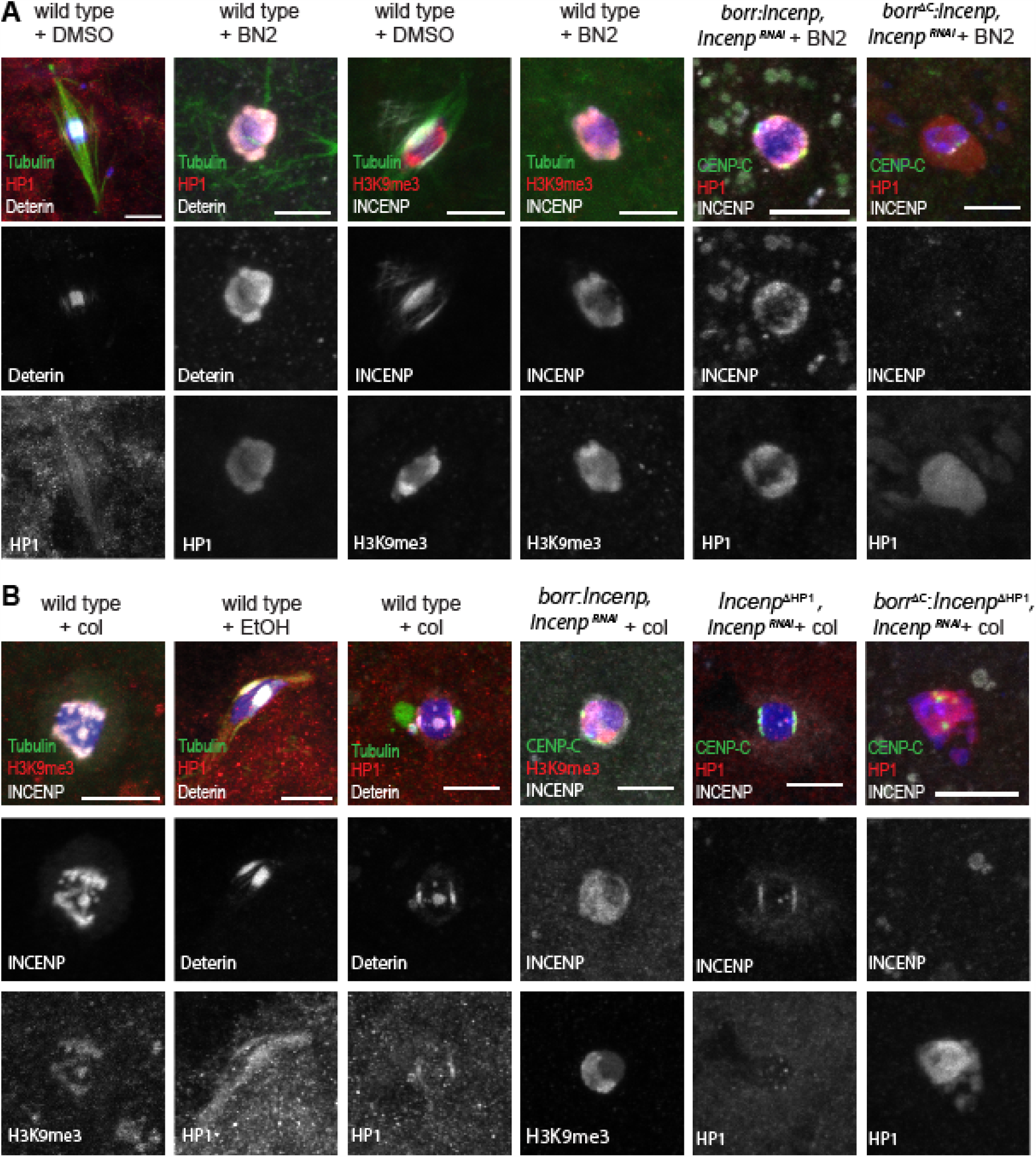
The dissociation of HP1 and the CPC from the chromosomes depends on microtubules. (A) Wild-type oocytes were treated with DMSO (left panel) or BN2 (right panel) for one hour to inhibit Aurora B kinase activity. In red are heterochromatic marks HP1 or H3K9me3 and in white are the CPC components INCENP or Deterin. In all the images, scale bars represent 5 µm. (B) Wild-type oocytes were treated for 60 minutes with ethanol or Colchicine to depolymerize microtubules and analyzed for the same markers as in (A). In all images, DNA is in blue and tubulin is in green.

In wild-type oocytes, HP1 is on the spindle but in the absence of Aurora B activity is on the chromosomes (Figure 5A). To test if Aurora B activity promotes the transfer of HP1 from the chromosomes to the spindle, we used colchicine to depolymerize the microtubules without inhibiting Aurora B activity. Colchicine treatment caused the CPC to retreat to the chromosomes and colocalize with H3K9me3 in wild-type oocytes (Figure 5B). Chromosome associated INCENP was also observed in colchicine-treated *borr:Incenp, Incenp* RNAi, consistent with the conclusion that Borealin promotes CPC localization to the chromosomes. However, HP1 was barely detectable in colchicine-treated wild-type oocytes (Figure 5B), unlike the observation in BN2-treated and *aurB* RNAi oocytes (Figure 5A and Figure 4A and B). These results suggested that Aurora B activity negatively regulates HP1 localization. To test this hypothesis, we compared colchicine treated *Incenp*^*ΔHP1*^, *Incenp* RNAi and *borr*^*ΔC*^*:Incenp*^*ΔHP1*^, *Incenp* RNAi oocytes. In the latter, which fail to recruit Aurora B to the chromosomes, HP1 localized to the chromatin. Because *Incenp*^*ΔHP1*^, *Incenp* RNAi oocytes recruit Aurora B to the chromosomes and assemble a spindle with associated HP1 while *borr*^*ΔC*^*:Incenp*^*ΔHP1*^, *Incenp* RNAi oocytes do not, the most likely explanation is that once HP1 is ejected from the chromosomes, Aurora B activity prevents its return. Thus, the ejection of HP1 from the chromosomes depends on Aurora B activity and Borealin. Ejection of the CPC from the chromosomes, however, depends on Aurora B activity, Borealin, and the microtubules.

### Recruitment of the CPC to the chromosomes and spindle assembly does not depend on INCENP microtubule interacting domains or Subito

We have thus far provided evidence that Borealin targets the CPC to the chromosomes, and then Aurora B activity results in spindle assembly and movement of the CPC to the microtubules. Because CPC localization to the chromosomes depends on the presence of microtubules, we examined if the microtubule binding domains within the CPC promoted spindle localization. INCENP has a single-α-helix (SAH) domain that binds microtubules (Samejima et al., 2015) and a conserved domain within the N-terminal region (the spindle transfer domain, STD) that is required for transfer to the midbody (Ainsztein et al., 1998), possibly by interacting with a kinesin 6 such as MKLP2/ Subito (Serena et al., 2020) (Figure S 1). To test if either of these microtubule interaction domains are important for the CPC to relocate and assemble the meiotic spindle, we generated deletions in each site (Figure 6A). In both *Incenp*^*ΔSTD*^, *Incenp* RNAi and *Incenp*^*ΔSAH*^, *Incenp* RNAi oocytes, we observed bipolar spindle assembly and normal CPC localization and a central spindle, suggesting these microtubule binding domains are not required for oocyte meiotic spindle assembly (Figure 6B). However, these females displayed either reduced fertility or sterility (Table 1), suggesting that these two domains of INCENP have an important role in embryonic mitosis. Whether these two domains are redundant in meiosis, or spindle localization of the CPC depends only on Borealin, or another pathway, to interact with microtubules, remains to be investigated.

**Fig 6:**
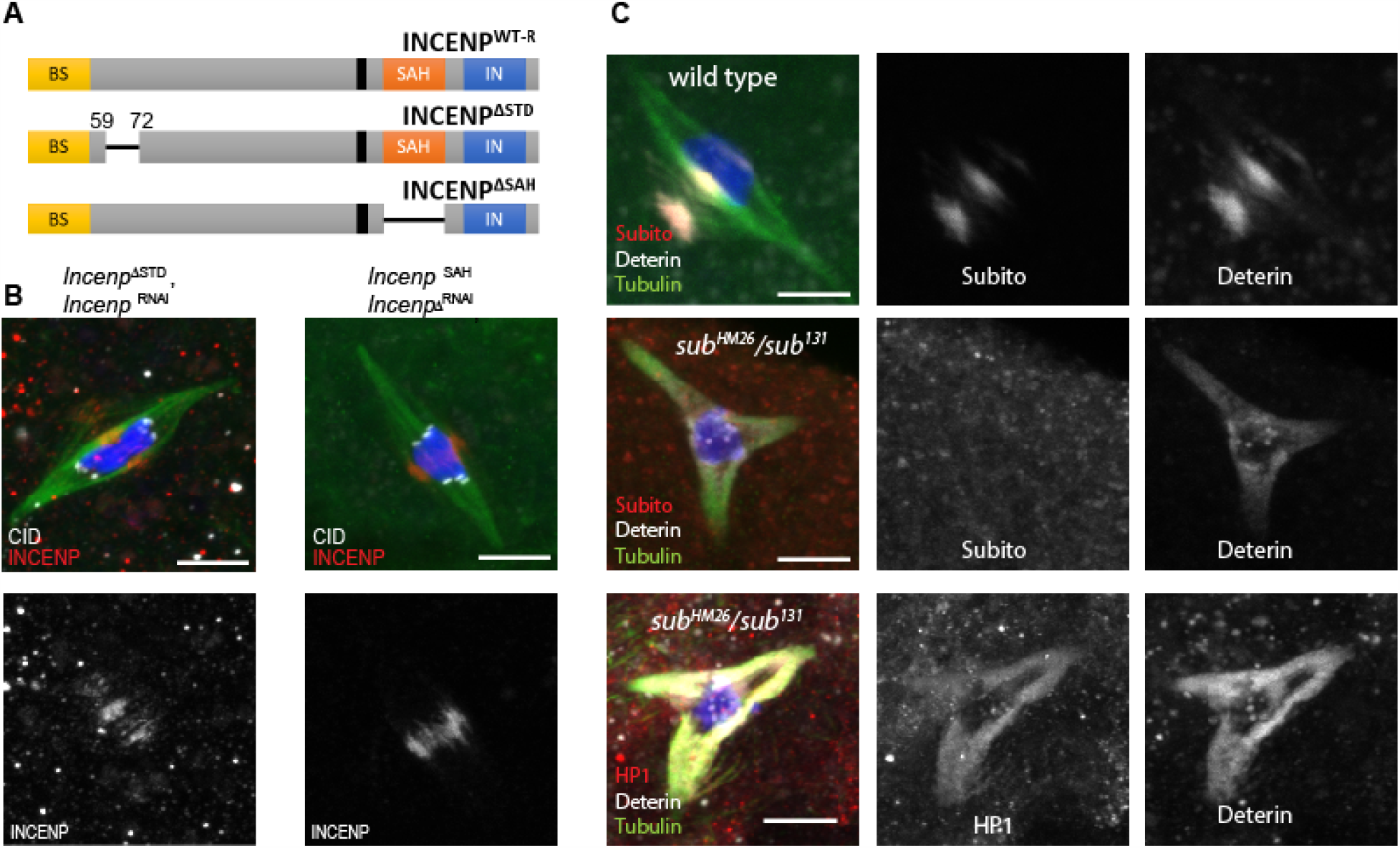
Analysis of CPC interactions with Subito and microtubules required for central spindle assembly. (A) A schematic showing two INCENP deletions removing regions that promote microtubule interactions. (B) Bipolar spindle assembly when *Incenp*^*ΔSTD*^ or *Incenp*^*ΔSAH*^ were expressed in *Incenp* RNAi oocytes. CID is in white, INCENP is in red, Tubulin is in green, DNA is in blue. (C) Tripolar spindle and mislocalization of the CPC phenotype in *sub*^*HM26*^/*sub*^*131*^ oocytes, with tubulin (green), Deterin (white), HP1 or Subito (red) localization. Scale bar is 5 µm in all images.

In *Drosophila*, kinesin-6 Subito is required to organize the central spindle, which includes recruiting the CPC (Das et al., 2018; Jang et al., 2005; Radford et al., 2012). However, the CPC can localize to the spindle microtubules in the absence of Subito (Das et al., 2018; Jang et al., 2005) (Figure 6C), suggesting that ejection of the CPC from the chromosomes may be sufficient for spindle transfer and Subito is not required. Interestingly, we found that Subito has a conserved HP1-binding site (amino acid 88-92, PQVFL). To test if the putative Subito HP1 binding site is required to build the central spindle, we examined *sub*^*HM26*^, a *sub* allele that has a point mutation (L92Q) in the HP1 binding site (Jang et al., 2005). Subito^HM26^ failed to localize to the spindle in oocytes, the spindle displayed a tripolar phenotype and the CPC localized throughout the spindle, all similar to a *sub* null mutant (Figure 6C). Thus, an HP1 interaction may be required for Subito localization, although we have not yet shown a direct interaction between HP1 and Subito.

### Homolog bi-orientation is regulated through the central spindle and proper spindle localization of the CPC and HP1

In mitotic cells, the CPC has an important role in error correction by destabilizing incorrect KT-MT attachments at the centromeres (Carmena et al., 2012b; Funabiki, 2019). Two hypomorphic CPC mutants, *Incenp*^*QA26*^ and *aurB*^*1689*^, could be defective in error correction because they are competent to build a bipolar spindle but have bi-orientation defects in oocytes (Radford et al., 2012; Resnick et al., 2009). Because the CPC in oocytes is most prominent on the central spindle, Aurora B activity could regulate bi-orientation while located on the microtubules rather than the chromosomes. To compare the role of CPC at the centromeres and central spindle in regulating homolog bi-orientation, we used fluorescence in situ hybridization (FISH) to examine three sets of *Incenp* mutants where the CPC is inappropriately localized to either the centromeres or the central spindle. The FISH probes targeted the pericentromeric regions each of chromosomes X, 2 and 3.

We first examined a sterile hypomorphic *Incenp* allele, *Incenp*^*18*.*197*^, which was discovered based on genetic interactions with *subito* (Das et al., 2016). In *Incenp*^*18*.*197*^ oocytes, the spindle was moderately diminished and a portion of INCENP was retained on the chromosomes in 67% of the oocytes (n=15) (Figure 7A), suggesting this mutant has a defect in CPC spindle transfer rather than chromosome localization. Using FISH, we determined that *Incenp*^*18*.*197*^ mutant oocytes have homolog bi-orientation defects (11%, n = 42). Second, *Incenp* transgenes with a MYC tag fused to the N-terminus have dominant defects in meiosis (Radford et al., 2012). To investigate whether the N-terminal tag was the cause of this defect, a new set of transgenes were constructed using the RNAi resistant backbone and different tags (MYC, HA and FLAG). We found that regardless of the fused tag, all the transgenes had similar phenotypes: reduced fertility and elevated meiotic nondisjunction in *Incenp* RNAi oocytes (Table 1), and failure to concentrate the CPC to the central spindle (Figure 7A). When the epitope tag was removed to generate *Incenp*^*WT-R*^ (this transgene was the backbone used to generate the mutants in this study), the defects in *Incenp* RNAi oocytes were fully restored to wild-type levels. These results confirmed that the N-terminal epitope tags in INCENP interfered with the CPC’s central spindle localization and function.

**Fig 7:**
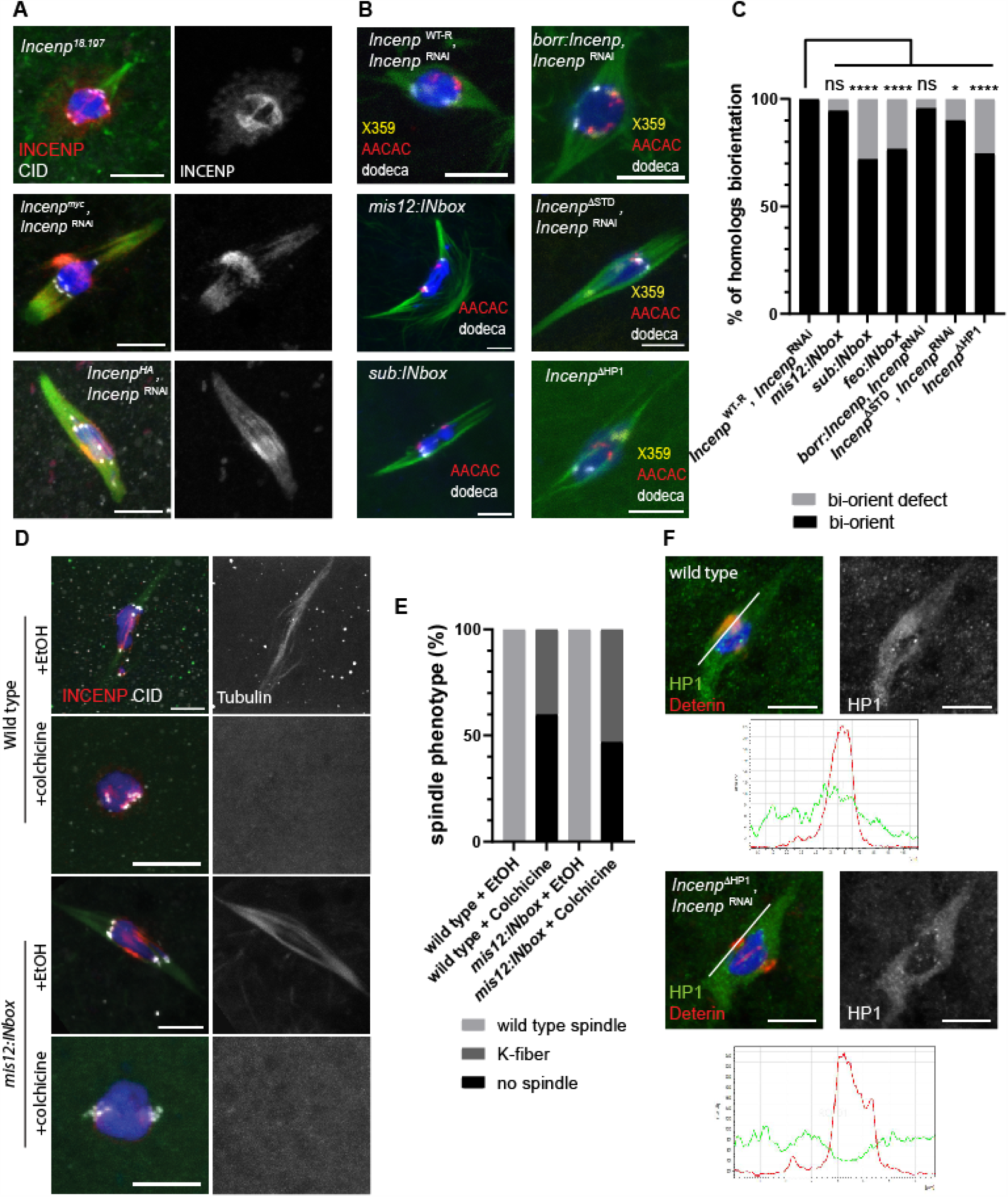
Disruption of homolog bi-orientation by disruptions of central spindle CPC. (A) Disorganized or mislocalized CPC in the *Incenp* hypomorphic allele *Incenp*^*18*.*197*^ and the transgenes *myc:Incenp and HA:Incenp*. The CPC or MYC are in red, CID is in white, tubulin is in green and DNA is in blue. (B) *Incenp* mutants examined for homolog bi-orientation using FISH with probes against pericentromeric heterochromatin on the X (359 bp repeat, yellow), 2^nd^ (AACAC, red) and 3^rd^ (dodeca, white) chromosomes. (C) Rates of bi-orientation defects were quantified (n= 57, 37, 50, 30, 69, 60 and 63 in the order of the graph). * = *p-*value < 0.05 ****= *p-*value <0.0001 in Fisher’s exact test. (D) Wild type oocytes and *mis12:INbox* oocytes treated with colchicine for 30 minutes. INCENP is in red and CID is in white. (E) Quantitation of spindle assembly after colchicine treatment (n= 5, 10, 7 and 17 in the order of the graph). (F) Localization of HP1 and Deterin in wild-type and *Incenp*^*ΔHP1*^, *Incenp* RNAi oocytes. HP1 is in green, Deterin is in red and overlapping region is in yellow.

Third, to test which population of the CPC regulates homolog bi-orientation, we used INbox fusions to target overexpression of Aurora B to specific sites. We predicted that overexpression of Aurora B would disrupt bi-orientation by destabilizing KT-MT attachments. Although forcing Aurora B localization to the centromere has been shown to cause bi-orientation defects in mitotic cells (Liu et al., 2009), the frequency of bi-orientation in oocytes expressing *mis12:INbox* was not significantly elevated compared to wild-type (Figure 7B and C). We also tested whether centromere-targeting Aurora B can destabilize microtubules by treating the oocytes with colchicine. K-fibers are more resistant to colchicine treatment than the central spindle, and the amount of K-fibers after colchicine treatment is a measure of attachment stability. The results of colchicine treated wild-type and *mis12:INbox* oocytes were comparable, the spindle was diminished to the same extent, indicating that the stability of the KT-MTs was similar in each genotype (Figure 7D and E). Thus, overexpression of Aurora B activity at the centromeres did not cause bi-orientation defects. In contrast, central-spindle-targeted Aurora B (*sub:INbox* or *feo:INbox*) caused significantly more bi-orientation defects than in wild type (Figure 7B and C). These results suggest that the CPC regulates homolog bi-orientation from within the central spindle rather than at the kinetochores.

If targeting the CPC to the central spindle is required for bi-orientation, *Incenp* mutants with defects interacting with microtubules should have bi-orientation defects. Indeed, *Incenp*^*ΔSTD*^, *Incenp* RNAi oocytes had homolog bi-orientation defects (Figure 7C), despite having apparently wild-type spindle assembly and CPC localization (Figure 5D). *Incenp*^*ΔHP1*^, *Incenp* RNAi oocytes had wild-type spindle morphology, but had defects in fertility and CPC localization to the central spindle (Figure 4D and G). Interestingly, we found HP1 spindle localization in *Incenp*^*ΔHP1*^, *Incenp* RNAi oocytes was different from wild type: HP1 was not enriched in the overlap with the CPC (Figure 7F). Like several of the *Incenp* mutants, *Incenp*^*ΔHP1*^ causes a dominant sterile phenotype (Table 1). Therefore, we examined *Incenp*^*ΔHP1*^ expressing oocytes by FISH and found these also had a homolog bi-orientation defect (Figure 7B and C). These results are consistent with the model that spindle localization of the CPC and interaction with HP1 is important for regulating homolog bi-orientation.

To test HP1 directly, we examined oocytes depleted of HP1 by expressing shRNA *GL00531* because null mutations in HP1 (*Su(var)205* in *Drosophila*), cause lethality. These oocytes displayed wild-type spindles with normal CPC localization (Figure S 4), which could be explained by the relatively mild knockdown of *HP1* (48% of mRNA remains). However, expression of *GL00531* in oocytes caused elevated X-chromosome non-disjunction (8.7%; n = 321). These results support the conclusion that during prometaphase I, HP1 and the CPC relocate from the chromosomes to the central spindle where they are both critical for homolog bi-orientation.

## Discussion

*Drosophila* oocytes generate spindles despite lacking centrioles or predefined spindle poles, like in many other organisms. Whether the microtubules assemble around MTOCs as in the mouse (Schuh and Ellenberg, 2007) or more closely around the chromosomes in *Drosophila* and humans (Hadders et al., 2020; Hengeveld et al., 2017), the chromatin appears to play a role in focusing microtubule assembly in the vicinity of, if not contacting, the chromosomes. The chromosome-associated molecules that drive this process, however, are not well known. Our previous studies have shown that the CPC is required for spindle assembly in *Drosophila* oocytes (Radford et al., 2012). We found that the known pathways for recruiting Survivin and Borealin to the centromere via Haspin and Bub1 are not essential for spindle assembly in *Drosophila* oocytes, although we have not ruled out minor roles in spindle assembly. Instead, we found that Borealin targets the CPC to chromatin, consistent with studies in *Xenopus* extracts (Kelly et al., 2007).

### CPC-dependent, chromosome-directed spindle assembly in oocytes depends on a Borealin-chromatin interaction, possibly involving HP1

Although INCENP can recruit HP1 (Kang et al., 2011), several lines of evidence suggest that HP1 can recruit the CPC prior to spindle assembly. When Aurora B is absent or inhibited in *Drosophila* oocytes, a complex of INCENP, Borealin and Survivin colocalizes with HP1 on chromosomes (Figure 5A). HP1 has also been shown to physically interact with the CPC in *Drosophila* (Alekseyenko et al., 2014). In HeLa cells, HP1 promotes CPC localization to chromatin and precedes H3T3 phosphorylation by Haspin kinase (Ruppert et al., 2018). Also in human cells, an interaction between Borealin and nucleosomes (Abad et al., 2019) or HP1 (Liu et al., 2014) recruits the CPC to the chromosomes. Borealin is also sufficient to recruit the CPC to chromatin in *Xenopus* (Kelly et al., 2007). Thus, the evidence from *Drosophila* oocytes and vertebrate cells is consistent and suggests that Borealin interacts with histones and HP1 to recruit the CPC to the chromatin. Possibly aided by HP1 interacting proteins like POGZ (Nozawa et al., 2010), phosphorylation of HP1 (Williams et al., 2019) or H3S10 (Duan et al., 2008; Fischle et al., 2005; Hirota et al., 2005) by Aurora B can promote HP1 ejection from the chromosomes. Thus, Aurora B activity could promote the transfer of a CPC-HP1 complex from the oocyte chromosomes to the microtubules.

We propose a model that not only explains how the chromosomes recruit spindle assembly factors, but also how the CPC moves from the chromosomes to the spindle (Figure 8). After nuclear envelope breakdown, a tripartite complex of the CPC comprised of INCENP, Borealin and Survivin/Deterin (Jeyaprakash et al., 2007) is recruited to the chromatin and enriched in regions marked by H3K9me3 and HP1. This localization is independent of Aurora B activity, suggesting that Borealin, in association with INCENP, is responsible for the recruitment of the CPC to HP1 and chromatin. Once the CPC localizes to the chromosomes, Aurora B activity results in phosphorylation of several targets and assembly of the kinetochore. HP1 is then ejected from the chromatin (Figure 4, Figure 5), which could be a mechanism for how the CPC is released from the chromatin and relocates onto the microtubules.

**Fig 8:**
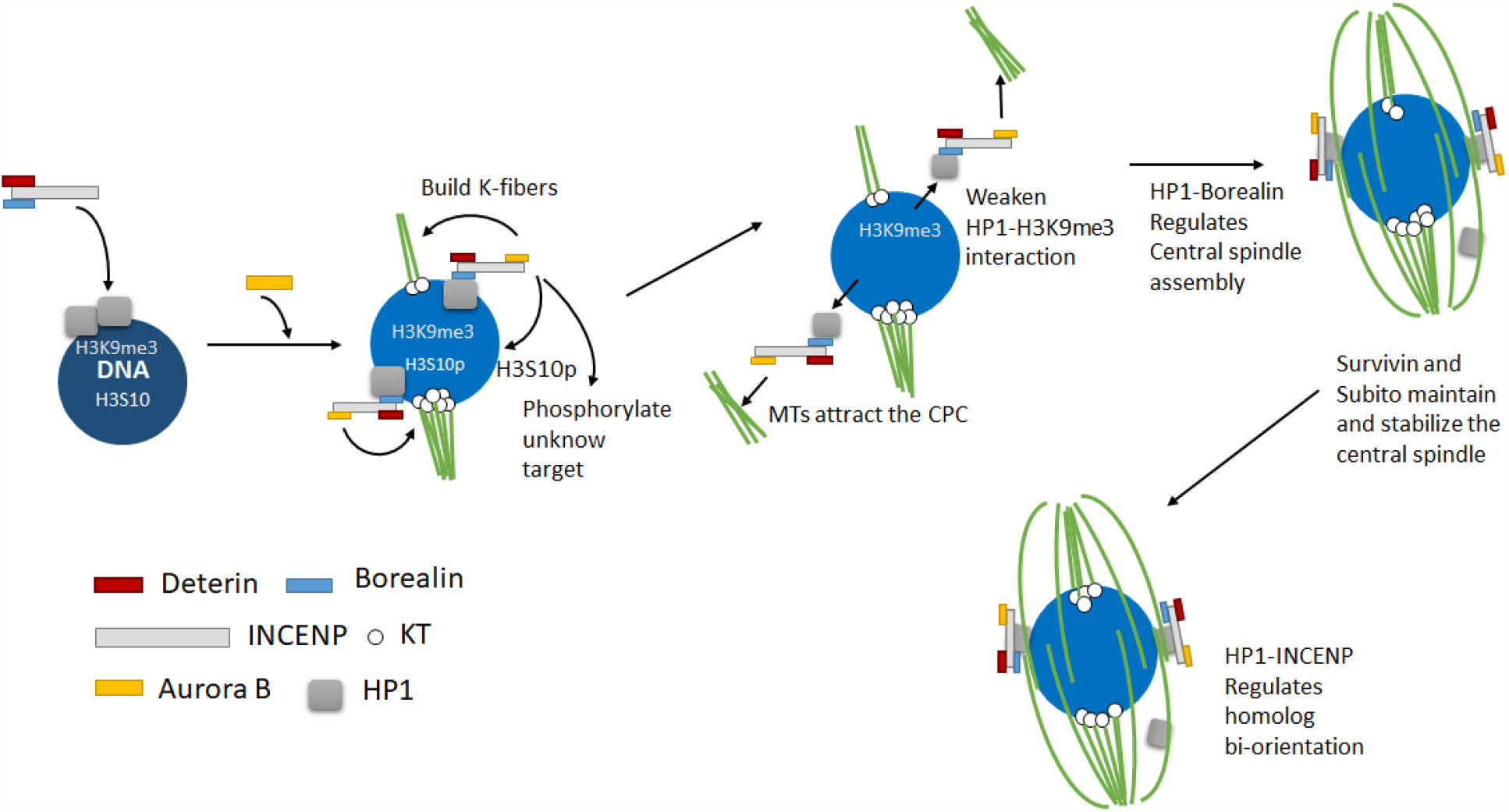
Model for spindle assembly in *Drosophila* oocytes. After Nuclear Envelope breakdown, a complex of INCENP, Borealin and Deterin / Survivin is recruited to the chromosomes. Localization studies suggest that CPC recruitment is enriched in heterochromatic regions containing H3K9me3 and HP1. Aurora B is recruited, which results in kinetochore assembly, limited microtubule recruitment in the form of K-fibers, and phosphorylation of other targets including H3S10 and possibly HP1. Aurora B activity also results in Borealin-dependent ejection of HP1 and the CPC from the chromosomes to the microtubules, although the target that is phosphorylated is not known. Once on the microtubules, the Kinesin 6 Subito causes enrichment of the CPC and HP1 in the central spindle.

This model explains how spindle assembly is restricted to the chromosomes in an acentrosomal system (Ohkura, 2015; Reschen et al., 2012; Rome and Ohkura, 2018); it is based on a Chromatin/HP1-Borealin interaction. We and others have suggested that restricting spindle assembly proximal to the chromosomes involves inhibition of spindle assembly factors in the cytoplasm (Beaven et al., 2017; Das et al., 2018; Rome and Ohkura, 2018). Spindle assembly could involve activating spindle promoting factors such as kinetochore proteins (Emanuele et al., 2008; Haase et al., 2017) and kinesins that bundle microtubules (Beaven et al., 2017; Das et al., 2018). For example, the kinesin NCD that has been shown to be inhibited by 14-3-3, which is released by Aurora B phosphorylation (Beaven et al., 2017). Spindle assembly may also involve suppressing microtubule depolymerases, such as kinesin 13/MCAK and Op18/Stathmin (Kelly et al., 2007; Sampath et al., 2004). We propose that the release of the CPC from the chromosomes locally activates spindle assembly factors.

### Non-kinetochore microtubules require spindle-associated CPC

In mutants where the CPC was targeted to the chromosomes (*mis12:Incenp, Det:Incenp)*, kinetochore assembly and K-fiber formation were observed (Figure 2, Figure 3). In other mutants (*HP1:Incenp* and *Incenp*^*ΔCEN*^), only limited kinetochore assembly and K-fibers were observed. Thus, low levels of CPC are sufficient for kinetochore assembly, but higher levels and/or specific localization are required for spindle assembly. Similar conclusions regarding localization and dosage have been made in *Xenopus;* kinetochore assembly can occur without localization of the CPC to the centromeres, may require less Aurora B activity than spindle assembly, and centromeric CPC localization is required for error correction but not for kinetochore assembly (Haase et al., 2017; Kelly et al., 2007; Tseng et al., 2010; Xu et al., 2009). A notable difference compared to *Xenopus*, however, is that the SAH domain is not required for kinetochore assembly in *Drosophila* oocytes (Bonner et al., 2019; Wheelock et al., 2017).

The absence of non-kinetochore microtubules in the mutants where the CPC was targeted to the chromosomes suggests spindle assembly requires microtubule associated CPC. For example, the DET:INCENP fusion promotes kinetochore and K-fiber assembly, but no central spindle (Figure 3). Central spindle assembly depends on transfer of the CPC from the chromosomes to the microtubules (Figure 5), and Deterin does not have this activity. The CPC contains multiple spindle-interacting domains, including two in INCENP (STD and SAH) (van der Horst et al., 2015). In addition, it has been proposed that an HP1-INCENP interaction in HeLa cells promotes the transfer of the CPC from the heterochromatin to the spindle (Ainsztein et al., 1998). However, it is possible that Borealin provides this activity in *Drosophila* oocytes. Borealin has a microtubule binding site (Trivedi et al., 2019b), which could drive spindle transfer, and explain how the BORR:INCENP fusion is sufficient for oocyte spindle assembly but the DET:INCENP fusion is not. Deterin has a role in stabilizing the central spindle (Figure 3, Figure 8)

Subito is required to promote release of CPC from chromatin in *Drosophila* (Cesario et al., 2006) and human (Serena et al., 2020) mitotic cells, but this is not the case in oocytes. In *sub* mutants, the central spindle is absent but robust bundles of CPC-containing non-kinetochore microtubules form (Jang et al., 2005). These observations suggest the CPC promotes assembly and bundling of microtubules, independent of Subito. When the CPC is ejected from the chromosomes, it may activate the Augmin pathway that has been shown to increase the amount of spindle microtubules in *Drosophila* oocytes (Rome and Ohkura, 2018). Subito, like its human homolog (Adriaans et al., 2020), is required to transport or recruit the CPC to the central spindle. Preventing Subito from interacting with the CPC could be an important regulatory modification in oocytes to ensure that microtubules do not assemble in the absence of chromosomes (Figure 6).

### Regulation of Homolog Bi-orientation by the CPC

Several previous studies have suggested that chromosome localized CPC regulates error correction, bi-orientation, and checkpoint silencing (Andrews et al., 2004; Foley and Kapoor, 2013; Liu et al., 2009; Tanaka et al., 2002), although some of these functions may not require precise centromere localization (Hadders et al., 2020; Hengeveld et al., 2017). However, our analysis of multiple *Incenp* mutants suggests that chromosomal localization of the CPC may not promote these functions. For example, the centromere-targeting of the CPC in meiosis did not cause KT-MT destabilization or affect homolog bi-orientation (Figure 7), as might be predicted if centromere-bound CPC can promote destabilization of microtubule attachments. Instead, several lines of evidence show that mutants with defects specific to spindle localization had the most severe bi-orientation defects (Figure 7). First, we have previously shown that defects in central spindle associated proteins, including actin-associated factors, results in bi-orientation defects (Das et al., 2016). Second, forcing localization of CPC to the central spindle, but not the kinetochores, disrupts bi-orientation. Third, an N-terminal epitope-tag on INCENP caused the CPC to spread out along the spindle, causing defects in homolog bi-orientation and fertility (Radford et al., 2012). These effects are enhanced by a reduction in Subito activity (Das et al., 2016; Radford et al., 2012), consistent with a function for the CPC in the central spindle. Finally, INCENP^ΔSTD^ oocytes had defective homolog bi-orientation, suggesting the conserved spindle transfer domain in INCENP is required for homolog bi-orientation. All these results suggest that defects in the central spindle caused a homolog bi-orientation defect, and are consistent with the hypothesis that homolog bi-orientation of meiotic chromosomes depends on interactions between the CPC and microtubules of the central spindle.

An INCENP-HP1 interaction also appear to be important for bi-orientation once the CPC and HP1 moves onto the spindle. Indeed, deleting the HP1 interaction site (121-232 amino acid) of INCENP caused disorganized CPC central spindle localization, loss of HP1 enrichment with the CPC and bi-orientation defects (Figure 4, Figure 7). HP1 or heterochromatin has also been shown to promote accurate achiasmate chromosome segregation during meiosis I in *Drosophila* oocytes (Giauque and Bickel, 2016; Karpen et al., 1996). HP1 interacts with a variety of proteins through its chromo-shadow domain (Eissenberg and Elgin, 2014). Therefore, HP1 could be involved in a complex pattern of interactions that bring important spindle proteins together, such as the CPC and Aurora B phosphorylation substrates such as Subito, which also has a conserved HP1 binding site that is required for its meiotic functions (Figure 7) (Jang et al., 2007; Jang et al., 2005). How this milieu of proteins promotes bi-orientation is still a mystery. The central spindle is a complex structure, containing several proteins that have a microtubule binding domains, including Borealin and INCENP, that may allow the CPC to simultaneously interact with microtubules and regulate KT-MT attachments (Trivedi et al., 2019b; Wheelock et al., 2017). Several *Drosophila* central spindle components have been suggested to form structures by phase separation (So et al., 2019), including HP1 (Liu et al., 2020) and the CPC (Trivedi et al., 2019a). We suggest the central spindle forms a unique structure that allows for the sensing bi-orientation of bivalents.

During meiosis I, the pairs of centromeres within a bivalent bi-orient at a much greater distance than the sister centromeres in mitosis or meiosis II. Therefore, it is plausible that the bi-orientation mechanism that works for sister centromeres does not work for homologous centromeres that are significantly further apart and lacking a direct connection. The meiotic central spindle may provide a direct connection between homologous centromeres by combining two properties. The first is a mechanism to coordinate the movement and separate the kinetochores of bivalents. This may be analogous to the activity of “bridging fibers” which can separate pairs of sister kinetochores in mitosis (Simunic and Tolic, 2016; Vukusic et al., 2017). In *C. elegans* meiosis, the central spindle separates homologs for chromosome segregation by microtubule pushing (Laband et al., 2017). The second is a mechanism for microtubule bound CPC to regulate KT-MT attachments and error-correction, which has been observed in several contexts (Fink et al., 2017; Funabiki, 2019; Pamula et al., 2019; Trivedi et al., 2019b). This combination of activities would facilitate allow the central spindle to facilitate reductional chromosome segregation at anaphase I.

## Methods and materials

### Generation of RNAi resistant INCENP

To engineer RNAi resistant transgenes, we obtained *Incenp* cDNA (RE52507) from *Drosophila* Genomic Resource Center and cloned it into the pENTR vector (Invitrogen, Carlsbad, CA). We used the Change-it Site-directed Mutagenesis kit (Affymetrix) to introduce 8 silent mutations in the region corresponding to amino acids 437-441, which is complementary to *Incenp* shRNA (GL00279). (Figure 1B and Figure S 1). The primers for the site-directed mutagenesis are: 5’-ATGAGCTTTTCAACCCACTCCTGCAGTCGCCCGTCAAGATGCGCGTGGAGGCGTTCGA −3’ and 5’-TCGAACGCCTCCACGCGCATCTTGACGGGCGACTGCAGGAGTGGGTTGAAAAGCTC ATG −3’. RNAi resistant INCENP constructs including *Incenp*^*myc*^, *Incenp*^*HA*^ and *Incenp*^*Flag*^ were inserted into the pPMW, pPHW or pPFW vectors that carry the UASp-promoter using the LR Clonase reaction (Gateway, Invitrogen). The construct was then injected in *w* embryos by Model System Injections (Durham, NC). Multiple transgenic lines were selected to balance in a *y w* background and crossed to *mata4-GAL-VP16* with/without *Incenp* RNAi for further testing. The transgenic lines on the 3^rd^ chromosome were chosen for generating a recombinant line with *Incenp* RNAi if the phenotype were comparable with the ones on the X or 2^nd^ chromosome. Expressing *Incenp*^myc^ in an *Incenp* RNAi background rescued spindle assembly and kinetochore assembly in oocytes as well as spindle localization. However, several defects were also observed, such as reduced fertility and elevated X-chromosome nondisjunction and the transgene protein was mislocalized along the spindle instead of concentrating in the central spindle. The same defects were observed previously with an *Incenp*^*myc*^ variant without the silent mutations (Radford et al., 2012). These results suggest that an epitope tag in the N-terminus of INCENP might interfere with its function, although the HA-tag may have less impact than the other epitopes (Table 1). To solve this problem, a Gibson Assembly kit (New England Biolabs) was used to remove the myc-tag from *Incenp*^*myc*^ to generate *Incenp*^*WT-R*^. Expressing *Incenp*^*WT-R*^ in *Incenp* RNAi oocytes displayed wild-type spindle and localization, and restored fertility to wild-type levels. We used a plasmid carrying *Incenp*^*WT-R*^ as the backbone for Gibson assembly reactions to generate all the *Incenp* mutations and fusions used in this study. For each mutation, at least two transgenic lines were analyzed for the ability to rescue *Incenp* RNAi with shRNA GL00279.

INbox constructs were generated by taking the last 101 amino acids (655-755) of INCENP including INbox and TSS activation site. Fusion proteins of INCENP were created by using MIS12 cDNA (RE19545), Deterin cDNA (LP03704), Su(var)205 cDNA (LD10408) and Borealin cDNA (LD36125). The constructs were injected into *Drosophila y w* embryos by Model System Injections (Durham, NC).

### *Drosophila* genetics and generation of shRNA transgenics

Flies were crossed and maintained on the standard media at 25°C. All loci information was obtained from Flybase. Fly stocks were obtained from the Bloomington Stock Center or the Transgenic RNAi Project at Harvard Medical School (TRiP, Boston, USA), including *aurB* (GL00202), *Incenp* (GL00279), *haspin* (GL00176), *Su(var)205* (GL00531) and *Bub1* (GL00151), except *mis12::EGFP* (Głuszek et al., 2015). To generate *Deterin* (LW501) and *haspin* (HK420) shRNA lines, a *Deterin* sequence (5’-CGGGAGAATGAGAAGCGTCTA −3’) or a *haspin* sequence (5’-GGAAGACAGTAGAGACAAATG-3’) were cloned into pVALIUM22 following the protocols described by the Harvard TRiP center. The construct was injected into *Drosophila* embryos (*y sc v; attP40*).

The pVALIUM22 vector carries the UASp promoter allowing for expression of short hairpins for RNA silencing and transgenes using the UAS/GAL4 binary expression system (Rorth et al., 1998). All the shRNA lines and transgenes used in this paper were expressed by *mata4-GAL-VP16*, which induces expression after early pachytene throughout most stages of oocyte development in *Drosophila* (Sugimura and Lilly, 2006).

For quantifying the knockdown of these RNAi lines, total RNA was extracted from late-stage oocytes using TRIzol® Reagent (Life Technologies) and reverse transcribed into cDNA using the High Capacity cDNA Reverse Transcription Kit (Applied Biosystems). The qPCR was performed on a StepOnePlus™ (Life Technologies) real-time PCR system using TaqMan® Gene Expression Assays (Life Technologies). Dm03420510_g1 for *haspin*, Dm02141491_g1 for *Deterin*, Dm01804657_g1 for *Bub1*, Dm0103608_g1 for *Su(var)205* and Dm02134593_g1 for the control *RpII140*. The knockdown of the respective mRNAs in these oocytes was reduced to 15% in *haspin* HK420 RNAi oocytes, 32% in *haspin* GL00176 RNAi oocytes, 5% in *Deterin* LW501 RNAi oocytes, 2% in *Bub1* GL00151 RNAi oocytes and 48% in *Su(var)205* GL00531 RNAi oocytes. To test for effects on mitosis, shRNA lines were tested for lethality when under the control of *P{tubP-GAL4}LL7*, which results in ubiquitous expression.

### Antibodies and immunofluorescence microscopy

To collect images of the meiotic spindle, oocytes were collected from 100-200, 3-4 day old yeast-fed non-virgin females. The protocol for fixation and immunofluorescence of stage 14 oocytes has been described (Radford and McKim, 2016). To observe whether spindle assembly was affected in post-meiotic mitosis, embryos were collected from several hundreds of yeast-fed females for 2 hours. After removing chorion by treated embryos with 50% bleach for 90 seconds, embryos then moved to tubes containing 500 µL heptane and 500 µL methanol and shaken vigorously for 30 seconds to fix. Rehydrated embryos were processed for immunofluorescence microscopy. Hoechst 33342 (10 µg/ml, Invitrogen) was used for staining DNA and mouse anti-a tubulin monoclonal antibody DM1A (1:50) conjugated with FITC (Sigma, St. Louis) was used for staining microtubules. Primary antibodies used in this paper were rabbit anti-CID (1:1000, Active motif), rabbit anti-SPC105R (1:4000, (Schittenhelm et al., 2007)), rabbit anti-CENP-C (1:5000, (Heeger et al., 2005)), mouse anti-Myc (1:50, 9E10, Roche, Indianapolis), mouse anti-Flag (1:500, Thermo Fisher), rat anti-INCENP (1:400, (Wu et al., 2008)), rabbit anti-Aurora B (1:1000, (Giet and Glover, 2001)), rabbit anti-Survivin (1:1000, (Szafer-Glusman et al., 2011)), rabbit anti-Borealin (1:100, (Gao et al., 2008)), mouse anti-HP1 (1:50, C1A9, Developmental Hybridoma Bank), rabbit anti-H3K9me3 (1:1000, Active motif), rat anti-Subito (1:75, (Jang et al., 2005)), rat anti-a-tubulin (Clone YOL 1/34, Millipore) and rabbit anti-pINCENP (1:1000, (Salimian et al., 2011)). The secondary antibodies including Cy3 and AlexFluor647 (Jackson Immunoresearch West Grove, PA) or AlexFluor488 (Molecular Probes) were used in accordance with the subjected primary antibodies. FISH probes for the X-chromosome (359 repeats), 2nd chromosome (AACAC satellite) and 3rd chromosome (dodeca satellite) and conjugated to either Alexa Flour 594, Cy3 or Cy5 were obtained from Integrated DNA Technologies (Dernburg et al., 1996; Radford and McKim, 2016). Oocytes were mounted in SlowFade Gold (Invitrogen). Images were collected on a Leica TCS SP8 confocal microscope with a 63x, NA 1.4 lens and shown as maximum projections of complete image stacks. Images were then cropped in Adobe Photoshop.

### Drug treatment assays

To inhibit Aurora B Kinase activity, oocytes were treated by either 0.1% DMSO or 50 µM BN2 in 0.1% DMSO in 60 minutes before fixation in Robb’s media. To depolymerize microtubules, oocytes were incubated in 250 µM colchicine in 0.5% ethanol or only 0.5% ethanol as a control for either 30 or 60 minutes before fixation depends on whether we wanted to destabilize spindle microtubules (Figure 7) or completely remove all spindle microtubules (Figure 1 and Figure 5).

### X-chromosome nondisjunction assays

To determine whether each *Incenp* mutant transgenes affected meiotic chromosome segregation, we measured fertility and X-chromosome nondisjunction. Transgenic virgin females were generated by crossing *mata4-GAL-VP16* to either the *Incenp* transgene or the *Incenp* transgene with *Incenp* RNAi or other RNAi lines. These transgenic females were crossed to *y Hw w/B*^*S*^*Y* males. The males carry a dominant mutation, *Bar*, on the Y chromosome which makes chromosome mis-segregation phenotypically distinguishable in the progeny. Crosses were set in vials and fertility was measured based on the progeny number. To compensate for the inviability of nullo-X and triplo-X progeny, the nondisjunction rate was calculated as 2*(XXY and XO progeny)/ total progeny, where total progeny was 2*(XXY and XO progeny)+ XX and XY progeny.

### Image analysis and statistics

To measure kinetochore localization, SPC105R foci were identified based on size and intensity and then counted using Imaris image analysis software (Bitplane) with the parameters used in Wang et. al. (Wang et al., 2019). To determine whether HP1 and the CPC were colocalized, line scans were drawn from pole to pole across the central spindle. The intensities of HP1 and Deterin were measured by using Leica SP8 software. In addition, the colocalizing region is shown in Figure 7F. When measuring bi-orientation by FISH, each data point corresponds to one pair of homologous chromosomes. Homologous chromosomes were considered bi-oriented if two FISH signals localized at the opposite ends of chromosome mass. Pairs of homologous chromosomes on the same side of the spindle were considered to have a bi-orientation defect. The appropriate statistical tests for each experiment as indicated in the figure legends were performed using Prism software (GraphPad).

## Acknowledgements

We thank Li Nguyen for technical assistance, Christian Lehner, James Wakefield and Michael Lampson for providing antibodies, and Karen Schindler, Ruth Steward for helpful comments and advice. We also thank Sarah Radford for early work on this project and comments on the manuscript. We thank the TRiP at Harvard Medical School for providing transgenic RNAi fly stocks used in this study. Fly stocks obtained from the Bloomington *Drosophila* Stock Center (NIH P40OD018537) were also used in this study. L.W. was funded by a Busch Predoctoral Fellowship. This work was supported by NIH grant GM101955 to K.S.M.

## Figure legends

Figure S 1: **Alignment and domain analysis of *Drosophila***.

The sequence alignment compares *D. melanogaster* INCENP to *D. virilis* and *X. laevis*. The Borealin/ Deterin binding domain is from amino acids 1-46 (yellow). The *CEN* and *STD* deletion mutations are marked red, two potential HP1 interaction sites are marked in blue, the RNAi mismatch region is marked in black, the SAH domain is marked in orange, and the INbox (IN) is in black.

Figure S 2: **Expression of *mis12:INbox* in the wild-type oocytes disrupts meiotic progression**.

(A) MIS12 localization in wild type and *Incenp* RNAi oocytes (arrows). MIS12 is in red, tubulin is in green, DNA is in blue, CENP-C is in white, and the scale bar represents 5 µm. (B) Fertilized 0-2hr old *Drosophila* embryos were fixed and stained for INCENP (red), tubulin (green) and DNA (blue). The scale bar is 5 µm. (C) Quantitation of SPC105R localization in oocytes with *mis12* fusions (n= 6, 9,12 oocytes). Error bars indicate 95% confidence intervals and ** = *p*-value < 0.01 run by Fisher’s exact test. (D) Expression of *Myc:feo:INbox* in wild-type and *mis12:INbox, Incenp*^RNAi^ oocytes. *Myc:feo:INbox* localizes to the central spindle in the wild-type oocytes. (E) Co-expression of *FLAG:mis12:INbox* and *Myc:feo:INbox* in *Incenp* RNAi oocytes. In these images, the Myc tag (FEO:INbox) is red, Aurora B or CID is white, tubulin is green and DNA is blue. The scale bar is 5 µm.

Figure S 3: **Borealin localization in *Det:Incenp, Incenp* RNAi oocytes and the localization of the CPC to the chromosmes**.

(A) Metaphase I oocytes from wild-type and *Det:Incenp, Incenp* RNAi females. Borealin is in white, INCENP is in red, tubulin is in green and DNA is in blue. Scale bar is 5 µm. (B) Expression of *Incenp* transgenes shown in (Figure 4C) in *Incenp* RNAi oocytes, including *borr*^*ΔC*^*:Incenp, Incenp*^*ΔHP1*^ and *borr*^*ΔC*^*:Incenp*^*ΔHP1*^. The images show SPC105R in white, INCENP in red, DNA in blue and tubulin in green. Scale bars represent 5 µm.

Figure S 4: **Phenotype of *haspin, Bub1* and *Su(var)205* (*HP1*) knockdowns**.

(A) Metaphase I oocytes from *haspin or Bub1 single RNAi* or *haspin, Bub1* double RNAi females. Centromere protein CID is in white, INCENP is in red, tubulin is in green and DNA is in blue. Scale bar is 5 µm. (B) Fertility and X-chromosome nondisjunction in *haspin* and *Bub1* RNAi females. G) Spindle in *Drosophila* HP1 mutant, *Su(var)205* (*HP1*) RNAi oocytes. CID is in white, INCENP is in red, tubulin is in green and DNA is in blue. Scale bars in all images are 5 µm.

## Literature cited

Abad, M.A., J.G. Ruppert, L. Buzuk, M. Wear, J. Zou, K.M. Webb, D.A. Kelly, P. Voigt, J. Rappsilber, W.C. Earnshaw, and A.A. Jeyaprakash. 2019. Borealin-nucleosome interaction secures chromosome association of the chromosomal passenger complex. J Cell Biol. 218:3912–3925.

Adams, R.R., H. Maiato, W.C. Earnshaw, and M. Carmena. 2001. Essential roles of Drosophila inner centromere protein (INCENP) and aurora B in histone H3 phosphorylation, metaphase chromosome alignment, kinetochore disjunction, and chromosome segregation. J Cell Biol. 153:865–880.

Adriaans, I.E., P.J. Hooikaas, A. Aher, M.J.M. Vromans, R.M. van Es, I. Grigoriev, A. Akhmanova, and S.M.A. Lens. 2020. MKLP2 Is a Motile Kinesin that Transports the Chromosomal Passenger Complex during Anaphase. Curr Biol. 30:2628–2637 e2629.

Ainsztein, A.M., S.E. Kandels-Lewis, A.M. Mackay, and W.C. Earnshaw. 1998. INCENP centromere and spindle targeting: identification of essential conserved motifs and involvement of heterochromatin protein HP1. J Cell Biol. 143:1763–1774.

Alekseyenko, A.A., A.A. Gorchakov, B.M. Zee, S.M. Fuchs, P.V. Kharchenko, and M.I. Kuroda. 2014. Heterochromatin-associated interactions of Drosophila HP1a with dADD1, HIPP1, and repetitive RNAs. Genes Dev. 28:1445–1460.

Andrews, P.D., Y. Ovechkina, N. Morrice, M. Wagenbach, K. Duncan, L. Wordeman, and J.R. Swedlow. 2004. Aurora B regulates MCAK at the mitotic centromere. Dev Cell. 6:253–268.

Beaven, R., R.N. Bastos, C. Spanos, P. Rome, C.F. Cullen, J. Rappsilber, R. Giet, G. Goshima, and H. Ohkura. 2017. 14-3-3 regulation of Ncd reveals a new mechanism for targeting proteins to the spindle in oocytes. J Cell Biol. 216:3029–3039.

Bennabi, I., M.E. Terret, and M.H. Verlhac. 2016. Meiotic spindle assembly and chromosome segregation in oocytes. J Cell Biol. 215:611–619.

Bishop, J.D., and J.M. Schumacher. 2002. Phosphorylation of the carboxyl terminus of inner centromere protein (INCENP) by the Aurora B Kinase stimulates Aurora B kinase activity. J Biol Chem. 277:27577–27580.

Bonner, M.K., J. Haase, J. Swinderman, H. Halas, L.M. Miller Jenkins, and A.E. Kelly. 2019. Enrichment of Aurora B kinase at the inner kinetochore controls outer kinetochore assembly. J Cell Biol. 218:3237–3257.

Carmena, M., X. Pinson, M. Platani, Z. Salloum, Z. Xu, A. Clark, F. Macisaac, H. Ogawa, U. Eggert, D.M. Glover, V. Archambault, and W.C. Earnshaw. 2012a. The chromosomal passenger complex activates Polo kinase at centromeres. PLoS Biol. 10:e1001250.

Carmena, M., M. Wheelock, H. Funabiki, and W.C. Earnshaw. 2012b. The chromosomal passenger complex (CPC): from easy rider to the godfather of mitosis. Nat Rev Mol Cell Biol. 13:789–803.

Cesario, J.M., J.K. Jang, B. Redding, N. Shah, T. Rahman, and K.S. McKim. 2006. Kinesin 6 family member Subito participates in mitotic spindle assembly and interacts with mitotic regulators. J Cell Sci. 119:4770–4780.

Chang, C.J., S. Goulding, R.R. Adams, W.C. Earnshaw, and M. Carmena. 2006. Drosophila Incenp is required for cytokinesis and asymmetric cell division during development of the nervous system. J Cell Sci. 119:1144–1153.

Cheeseman, I.M. 2014. The kinetochore. Cold Spring Harbor perspectives in biology. 6:a015826.

Colombié, N., C.F. Cullen, A.L. Brittle, J.K. Jang, W.C. Earnshaw, M. Carmena, K. McKim, and H. Ohkura. 2008. Dual roles of Incenp crucial to the assembly of the acentrosomal metaphase spindle in female meiosis. Development. 135:3239-3246.

Costa, M.0F.A., and H. Ohkura. 2019. The molecular architecture of the meiotic spindle is remodeled during metaphase arrest in oocytes. J Cell Biol. 218:2854–2864.

Das, A., J. Cesario, A.M. Hinman, J.K. Jang, and K.S. McKim. 2018. Kinesin 6 Regulation in Drosophila Female Meiosis by the Non-conserved N- and C-Terminal Domains. G3 (Bethesda, Md. 8:1555–1569.

Das, A., S.J. Shah, B. Fan, D. Paik, D.J. DiSanto, A.M. Hinman, J.M. Cesario, R.A. Battaglia, N. Demos, and K.S. McKim. 2016. Spindle Assembly and Chromosome Segregation Requires Central Spindle Proteins in Drosophila Oocytes. Genetics. 202:61–75.

Dernburg, A.F., J.W. Sedat, and R.S. Hawley. 1996. Direct evidence of a role for heterochromatin in meiotic chromosome segregation. Cell. 86:135–146.

Drutovic, D., X. Duan, R. Li, P. Kalab, and P. Solc. 2020. RanGTP and importin beta regulate meiosis I spindle assembly and function in mouse oocytes. EMBO J. 39:e101689.

Duan, Q., H. Chen, M. Costa, and W. Dai. 2008. Phosphorylation of H3S10 blocks the access of H3K9 by specific antibodies and histone methyltransferase. Implication in regulating chromatin dynamics and epigenetic inheritance during mitosis. J Biol Chem. 283:33585–33590.

Dumont, J., and A. Desai. 2012. Acentrosomal spindle assembly and chromosome segregation during oocyte meiosis. Trends Cell Biol. 22:241–249.

Dumont, J., K. Oegema, and A. Desai. 2010. A kinetochore-independent mechanism drives anaphase chromosome separation during acentrosomal meiosis. Nat Cell Biol. 12:894–901.

Eissenberg, J.C., and S.C. Elgin. 2014. HP1a: a structural chromosomal protein regulating transcription. Trends Genet. 30:103–110.

Emanuele, M.J., W. Lan, M. Jwa, S.A. Miller, C.S. Chan, and P.T. Stukenberg. 2008. Aurora B kinase and protein phosphatase 1 have opposing roles in modulating kinetochore assembly. J Cell Biol. 181:241–254.

Feijão, T., O. Afonso, A.F. Maia, and C.E. Sunkel. 2013. Stability of kinetochore-microtubule attachment and the role of different KMN network components in Drosophila. Cytoskeleton (Hoboken, N.J. 70:661–675.

Fink, S., K. Turnbull, A. Desai, and C.S. Campbell. 2017. An engineered minimal chromosomal passenger complex reveals a role for INCENP/Sli15 spindle association in chromosome biorientation. J Cell Biol. 216:911–923.

Fischle, W., B.S. Tseng, H.L. Dormann, B.M. Ueberheide, B.A. Garcia, J. Shabanowitz, D.F. Hunt, H. Funabiki, and C.D. Allis. 2005. Regulation of HP1-chromatin binding by histone H3 methylation and phosphorylation. Nature. 438:1116–1122.

Foley, E.A., and T.M. Kapoor. 2013. Microtubule attachment and spindle assembly checkpoint signalling at the kinetochore. Nat Rev Mol Cell Biol. 14:25–37.

Fresan, U., M.A. Rodriguez-Sanchez, O. Reina, V.G. Corces, and M.L. Espinas. 2020. Haspin kinase modulates nuclear architecture and Polycomb-dependent gene silencing. PLoS Genet. 16:e1008962.

Funabiki, H. 2019. Correcting aberrant kinetochore microtubule attachments: a hidden regulation of Aurora B on microtubules. Curr Opin Cell Biol. 58:34–41.

Gao, S., M.G. Giansanti, G.J. Buttrick, S. Ramasubramanyan, A. Auton, M. Gatti, and J.G. Wakefield. 2008. Australin: a chromosomal passenger protein required specifically for Drosophila melanogaster male meiosis. J Cell Biol. 180:521–535.

Giauque, C.C., and S.E. Bickel. 2016. Heterochromatin-Associated Proteins HP1a and Piwi Collaborate to Maintain the Association of Achiasmate Homologs in Drosophila Oocytes. Genetics. 203:173–189.

Giet, R., and D.M. Glover. 2001. Drosophila aurora B kinase is required for histone H3 phosphorylation and condensin recruitment during chromosome condensation and to organize the central spindle during cytokinesis. J Cell Biol. 152:669–682.

Giunta, K.L., J.K. Jang, E.A. Manheim, G. Subramanian, and K.S. McKim. 2002. subito encodes a kinesin-like protein required for meiotic spindle pole formation in Drosophila melanogaster. Genetics. 160:1489–1501.

Głuszek, A.A., C.F. Cullen, W. Li, R.A. Battaglia, S.J. Radford, M.F. Costa, K.S. McKim, G. Goshima, and H. Ohkura. 2015. The microtubule catastrophe promoter Sentin delays stable kinetochore-microtubule attachment in oocytes. J Cell Biol. 211:1113–1120.

Gohard, F.H., D.J. St-Cyr, M. Tyers, and W.C. Earnshaw. 2014. Targeting the INCENP IN-box-Aurora B interaction to inhibit CPC activity in vivo. Open Biol. 4:140163.

Haase, J., M.K. Bonner, H. Halas, and A.E. Kelly. 2017. Distinct Roles of the Chromosomal Passenger Complex in the Detection of and Response to Errors in Kinetochore-Microtubule Attachment. Dev Cell. 42:640–654 e645.

Hadders, M.A., S. Hindriksen, M.A. Truong, A.N. Mhaskar, J.P. Wopken, M.J.M. Vromans, and S.M.A. Lens. 2020. Untangling the contribution of Haspin and Bub1 to Aurora B function during mitosis. J Cell Biol. 219.

Heald, R., and A. Khodjakov. 2015. Thirty years of search and capture: The complex simplicity of mitotic spindle assembly. J Cell Biol. 211:1103–1111.

Heald, R., R. Tournebize, T. Blank, R. Sandaltzopoulos, P. Becker, A. Hyman, and E. Karsenti. 1996. Self-organization of microtubules into bipolar spindles around artificial chromosomes in Xenopus egg extracts. Nature. 382:420–425.

Heeger, S., O. Leismann, R. Schittenhelm, O. Schraidt, S. Heidmann, and C.F. Lehner. 2005. Genetic interactions of separase regulatory subunits reveal the diverged Drosophila Cenp-C homolog. Genes Dev. 19:2041–2053.

Hengeveld, R.C.C., M.J.M. Vromans, M. Vleugel, M.A. Hadders, and S.M.A. Lens. 2017. Inner centromere localization of the CPC maintains centromere cohesion and allows mitotic checkpoint silencing. Nat Commun. 8:15542.

Hindriksen, S., S.M.A. Lens, and M.A. Hadders. 2017. The Ins and Outs of Aurora B Inner Centromere Localization. Front Cell Dev Biol. 5:112.

Hirota, T., J.J. Lipp, B.H. Toh, and J.M. Peters. 2005. Histone H3 serine 10 phosphorylation by Aurora B causes HP1 dissociation from heterochromatin. Nature. 438:1176–1180.

Holubcová, Z., M. Blayney, K. Elder, and M. Schuh. 2015. Human oocytes. Error-prone chromosome-mediated spindle assembly favors chromosome segregation defects in human oocytes. Science. 348:1143–1147.

Jang, J.K., T. Rahman, V.S. Kober, J. Cesario, and K.S. McKim. 2007. Misregulation of the Kinesin-like Protein Subito Induces Meiotic Spindle Formation in the Absence of Chromosomes and Centrosomes. Genetics. 177:267–280.

Jang, J.K., T. Rahman, and K.S. McKim. 2005. The kinesinlike protein Subito contributes to central spindle assembly and organization of the meiotic spindle in Drosophila oocytes. Mol Biol Cell. 16:4684–4694.

Jeyaprakash, A.A., U.R. Klein, D. Lindner, J. Ebert, E.A. Nigg, and E. Conti. 2007. Structure of a Survivin-Borealin-INCENP core complex reveals how chromosomal passengers travel together. Cell. 131:271–285.

Kang, J., J. Chaudhary, H. Dong, S. Kim, C.A. Brautigam, and H. Yu. 2011. Mitotic centromeric targeting of HP1 and its binding to Sgo1 are dispensable for sister-chromatid cohesion in human cells. Mol Biol Cell. 22:1181–1190.

Karpen, G.H., M.H. Le, and H. Le. 1996. Centric heterochromatin and the efficiency of achiasmate disjunction in Drosophila female meiosis. Science. 273:118–122.

Kelly, A.E., S.C. Sampath, T.A. Maniar, E.M. Woo, B.T. Chait, and H. Funabiki. 2007. Chromosomal enrichment and activation of the aurora B pathway are coupled to spatially regulate spindle assembly. Dev Cell. 12:31–43.

Klein, U.R., E.A. Nigg, and U. Gruneberg. 2006. Centromere targeting of the chromosomal passenger complex requires a ternary subcomplex of Borealin, Survivin, and the N-terminal domain of INCENP. Mol Biol Cell. 17:2547–2558.

Krenn, V., and A. Musacchio. 2015. The Aurora B Kinase in Chromosome Bi-Orientation and Spindle Checkpoint Signaling. Front Oncol. 5:225.

Laband, K., R. Le Borgne, F. Edwards, M. Stefanutti, J.C. Canman, J.M. Verbavatz, and J. Dumont. 2017. Chromosome segregation occurs by microtubule pushing in oocytes. Nat Commun. 8:1499.

Liu, D., G. Vader, M.J. Vromans, M.A. Lampson, and S.M. Lens. 2009. Sensing chromosome bi-orientation by spatial separation of aurora B kinase from kinetochore substrates. Science. 323:1350–1353.

Liu, X., J. Shen, L. Xie, Z. Wei, C. Wong, Y. Li, X. Zheng, P. Li, and Y. Song. 2020. Mitotic Implantation of the Transcription Factor Prospero via Phase Separation Drives Terminal Neuronal Differentiation. Dev Cell. 52:277–293 e278.

Liu, X., Z. Song, Y. Huo, J. Zhang, T. Zhu, J. Wang, X. Zhao, F. Aikhionbare, J. Zhang, H. Duan, J. Wu, Z. Dou, Y. Shi, and X. Yao. 2014. Chromatin protein HP1 interacts with the mitotic regulator borealin protein and specifies the centromere localization of the chromosomal passenger complex. J Biol Chem. 289:20638–20649.

Maresca, T.J., A.C. Groen, J.C. Gatlin, R. Ohi, T.J. Mitchison, and E.D. Salmon. 2009. Spindle assembly in the absence of a RanGTP gradient requires localized CPC activity. Curr Biol. 19:1210–1215.

Matthies, H.J., H.B. McDonald, L.S. Goldstein, and W.E. Theurkauf. 1996. Anastral meiotic spindle morphogenesis: role of the non-claret disjunctional kinesin-like protein. J Cell Biol. 134:455–464.

Mullen, T.J., A.C. Davis-Roca, and S.M. Wignall. 2019. Spindle assembly and chromosome dynamics during oocyte meiosis. Curr Opin Cell Biol. 60:53–59.

Muscat, C.C., K.M. Torre-Santiago, M.V. Tran, J.A. Powers, and S.M. Wignall. 2015. Kinetochore-independent chromosome segregation driven by lateral microtubule bundles. eLife. 4.

Nicklas, R.B. 1997. How cells get the right chromosomes. Science. 275:632–637.

Nozawa, R.S., K. Nagao, H.T. Masuda, O. Iwasaki, T. Hirota, N. Nozaki, H. Kimura, and C. Obuse. 2010. Human POGZ modulates dissociation of HP1alpha from mitotic chromosome arms through Aurora B activation. Nat Cell Biol. 12:719–727.

Ohkura, H. 2015. Meiosis: an overview of key differences from mitosis. Cold Spring Harbor perspectives in biology. 7.

Pamula, M.C., L. Carlini, S. Forth, P. Verma, S. Suresh, W.R. Legant, A. Khodjakov, E. Betzig, and T.M. Kapoor. 2019. High-resolution imaging reveals how the spindle midzone impacts chromosome movement. J Cell Biol. 218:2529–2544.

Przewloka, M.R., W. Zhang, P. Costa, V. Archambault, P.P. D’Avino, K.S. Lilley, E.D. Laue, A.D. McAinsh, and D.M. Glover. 2007. Molecular analysis of core kinetochore composition and assembly in Drosophila melanogaster. PloS one. 2:e478.

Radford, S.J., T.L. Hoang, A.A. Głuszek, H. Ohkura, and K.S. McKim. 2015. Lateral and End-On Kinetochore Attachments Are Coordinated to Achieve Bi-orientation in Drosophila Oocytes. PLoS Genet. 11:e1005605.

Radford, S.J., J.K. Jang, and K.S. McKim. 2012. The Chromosomal Passenger Complex is required for Meiotic Acentrosomal Spindle Assembly and Chromosome Bi-orientation. Genetics. 192:417–429.

Radford, S.J., and K.S. McKim. 2016. Techniques for Imaging Prometaphase and Metaphase of Meiosis I in Fixed Drosophila Oocytes. J Vis Exp. 116:e54666.

Radford, S.J., A.L. Nguyen, K. Schindler, and K.S. McKim. 2017. The chromosomal basis of meiotic acentrosomal spindle assembly and function in oocytes. Chromosoma. 126:351–364.

Reschen, R.F., N. Colombie, L. Wheatley, J. Dobbelaere, D. St Johnston, H. Ohkura, and J.W. Raff. 2012. Dgp71WD is required for the assembly of the acentrosomal Meiosis I spindle, and is not a general targeting factor for the gamma-TuRC. Biol Open. 1:422–429.

Resnick, T.D., K.J. Dej, Y. Xiang, R.S. Hawley, C. Ahn, and T.L. Orr-Weaver. 2009. Mutations in the chromosomal passenger complex and the condensin complex differentially affect synaptonemal complex disassembly and metaphase I configuration in Drosophila female meiosis. Genetics. 181:875–887.

Rome, P., and H. Ohkura. 2018. A novel microtubule nucleation pathway for meiotic spindle assembly in oocytes. J Cell Biol. 217:3431–3445.

Rorth, P., K. Szabo, A. Bailey, T. Laverty, J. Rehm, G.M. Rubin, K. Weigmann, M. Milan, V. Benes, W. Ansorge, and S.M. Cohen. 1998. Systematic Gain-of-Function Genetics in Drosophila. Development. 125:1049–1057.

Ruppert, J.G., K. Samejima, M. Platani, O. Molina, H. Kimura, A.A. Jeyaprakash, S. Ohta, and W.C. Earnshaw. 2018. HP1alpha targets the chromosomal passenger complex for activation at heterochromatin before mitotic entry. EMBO J. 37.

Salimian, K.J., E.R. Ballister, E.M. Smoak, S. Wood, T. Panchenko, M.A. Lampson, and B.E. Black. 2011. Feedback control in sensing chromosome biorientation by the Aurora B kinase. Curr Biol. 21:1158–1165.

Samejima, K., M. Platani, M. Wolny, H. Ogawa, G. Vargiu, P.J. Knight, M. Peckham, and W.C. Earnshaw. 2015. The Inner Centromere Protein (INCENP) Coil Is a Single alpha-Helix (SAH) Domain That Binds Directly to Microtubules and Is Important for Chromosome Passenger Complex (CPC) Localization and Function in Mitosis. J Biol Chem. 290:21460–21472.

Sampath, S.C., R. Ohi, O. Leismann, A. Salic, A. Pozniakovski, and H. Funabiki. 2004. The chromosomal passenger complex is required for chromatin-induced microtubule stabilization and spindle assembly. Cell. 118:187–202.

Schittenhelm, R.B., S. Heeger, F. Althoff, A. Walter, S. Heidmann, K. Mechtler, and C.F. Lehner. 2007. Spatial organization of a ubiquitous eukaryotic kinetochore protein network in Drosophila chromosomes. Chromosoma. 116:385–402.

Schuh, M., and J. Ellenberg. 2007. Self-organization of MTOCs replaces centrosome function during acentrosomal spindle assembly in live mouse oocytes. Cell. 130:484–498.

Serena, M., R.N. Bastos, P.R. Elliott, and F.A. Barr. 2020. Molecular basis of MKLP2-dependent Aurora B transport from chromatin to the anaphase central spindle. J Cell Biol. 219.

Simunic, J., and I.M. Tolic. 2016. Mitotic Spindle Assembly: Building the Bridge between Sister K-Fibers. Trends Biochem Sci. 41:824–833.

Smurnyy, Y., A.V. Toms, G.R. Hickson, M.J. Eck, and U.S. Eggert. 2010. Binucleine 2, an isoform-specific inhibitor of Drosophila Aurora B kinase, provides insights into the mechanism of cytokinesis. ACS Chem Biol. 5:1015–1020.

So, C., K.B. Seres, A.M. Steyer, E. Monnich, D. Clift, A. Pejkovska, W. Mobius, and M. Schuh. 2019. A liquid-like spindle domain promotes acentrosomal spindle assembly in mammalian oocytes. Science. 364.

Sugimura, I., and M.A. Lilly. 2006. Bruno inhibits the expression of mitotic cyclins during the prophase I meiotic arrest of Drosophila oocytes. Dev Cell. 10:127–135.

Szafer-Glusman, E., M.T. Fuller, and M.G. Giansanti. 2011. Role of Survivin in cytokinesis revealed by a separation-of-function allele. Mol Biol Cell. 22:3779–3790.

Tanaka, T.U., N. Rachidi, C. Janke, G. Pereira, M. Galova, E. Schiebel, M.J. Stark, and K. Nasmyth. 2002. Evidence that the Ipl1-Sli15 (Aurora kinase-INCENP) complex promotes chromosome biorientation by altering kinetochore-spindle pole connections. Cell. 108:317–329.

Theurkauf, W.E., and R.S. Hawley. 1992. Meiotic spindle assembly in Drosophila females: behavior of nonexchange chromosomes and the effects of mutations in the nod kinesin-like protein. J Cell Biol. 116:1167–1180.

Trivedi, P., F. Palomba, E. Niedzialkowska, M.A. Digman, E. Gratton, and P.T. Stukenberg. 2019a. The inner centromere is a biomolecular condensate scaffolded by the chromosomal passenger complex. Nat Cell Biol. 21:1127–1137.

Trivedi, P., and P.T. Stukenberg. 2020. A Condensed View of the Chromosome Passenger Complex. Trends Cell Biol.

Trivedi, P., A.V. Zaytsev, M. Godzi, F.I. Ataullakhanov, E.L. Grishchuk, and P.T. Stukenberg. 2019b. The binding of Borealin to microtubules underlies a tension independent kinetochore-microtubule error correction pathway. Nat Commun. 10:682.

Tseng, B.S., L. Tan, T.M. Kapoor, and H. Funabiki. 2010. Dual Detection of Chromosomes and Microtubules by the Chromosomal Passenger Complex Drives Spindle Assembly. Dev Cell. 18:903–912.

van der Horst, A., and S.M. Lens. 2014. Cell division: control of the chromosomal passenger complex in time and space. Chromosoma. 123:25–42.

van der Horst, A., M.J. Vromans, K. Bouwman, M.S. van der Waal, M.A. Hadders, and S.M. Lens. 2015. Inter-domain Cooperation in INCENP Promotes Aurora B Relocation from Centromeres to Microtubules. Cell reports. 12:380–387.

Venkei, Z., M.R. Przewloka, Y. Ladak, S. Albadri, A. Sossick, G. Juhasz, B. Novak, and D.M. Glover. 2012. Spatiotemporal dynamics of Spc105 regulates the assembly of the Drosophila kinetochore. Open Biol. 2:110032.

Vukusic, K., R. Buda, A. Bosilj, A. Milas, N. Pavin, and I.M. Tolic. 2017. Microtubule Sliding within the Bridging Fiber Pushes Kinetochore Fibers Apart to Segregate Chromosomes. Dev Cell. 43:11–23 e16.

Wang, F., J. Dai, J.R. Daum, E. Niedzialkowska, B. Banerjee, P.T. Stukenberg, G.J. Gorbsky, and J.M. Higgins. 2010. Histone H3 Thr-3 phosphorylation by Haspin positions Aurora B at centromeres in mitosis. Science. 330:231–235.

Wang, L.I., A. Das, and K.S. McKim. 2019. Sister centromere fusion during meiosis I depends on maintaining cohesins and destabilizing microtubule attachments. PLoS Genet. 15:e1008072.

Watanabe, Y. 2012. Geometry and force behind kinetochore orientation: lessons from meiosis. Nat Rev Mol Cell Biol. 13:370–382.

Wheelock, M.S., D.J. Wynne, B.S. Tseng, and H. Funabiki. 2017. Dual recognition of chromatin and microtubules by INCENP is important for mitotic progression. J Cell Biol. 216:925–941.

Wignall, S.M., and A.M. Villeneuve. 2009. Lateral microtubule bundles promote chromosome alignment during acentrosomal oocyte meiosis. Nat Cell Biol. 11:839–844.

Williams, M.M., A.J. Mathison, T. Christensen, P.T. Greipp, D.L. Knutson, E.W. Klee, M.T. Zimmermann, J. Iovanna, G.A. Lomberk, and R.A. Urrutia. 2019. Aurora kinase B-phosphorylated HP1alpha functions in chromosomal instability. Cell Cycle. 18:1407–1421.

Wu, C., V. Singaram, and K.S. McKim. 2008. mei-38 is required for chromosome segregation during meiosis in Drosophila females. Genetics. 180:61–72.

Xu, Z., H. Ogawa, P. Vagnarelli, J.H. Bergmann, D.F. Hudson, S. Ruchaud, T. Fukagawa, W.C. Earnshaw, and K. Samejima. 2009. INCENP-aurora B interactions modulate kinase activity and chromosome passenger complex localization. J Cell Biol. 187:637–653.

Yamagishi, Y., T. Honda, Y. Tanno, and Y. Watanabe. 2010. Two histone marks establish the inner centromere and chromosome bi-orientation. Science. 330:239–243.

