## Supplemental Figures for "Oocyte spindle assembly depends on multiple interactions between HP1 and the CPC"

Figure S 4: **Phenotype of *haspin, Bub1* and *Su(var)205* (*HP1*) knockdowns.**

(A) Metaphase I oocytes from *haspin or Bub1 single RNAi* or *haspin, Bub1* double RNAi females. Centromere protein CID is in white, INCENP is in red, tubulin is in green and DNA is in blue. Scale bar is 5 µm. (B) Fertility and X-chromosome nondisjunction in *haspin* and *Bub1* RNAi females. G) Spindle in *Drosophila* HP1 mutant, *Su(var)205* (*HP1*) RNAi oocytes. CID is in white, INCENP is in red, tubulin is in green and DNA is in blue. Scale bars in all images are 5 µm.

Figure S1

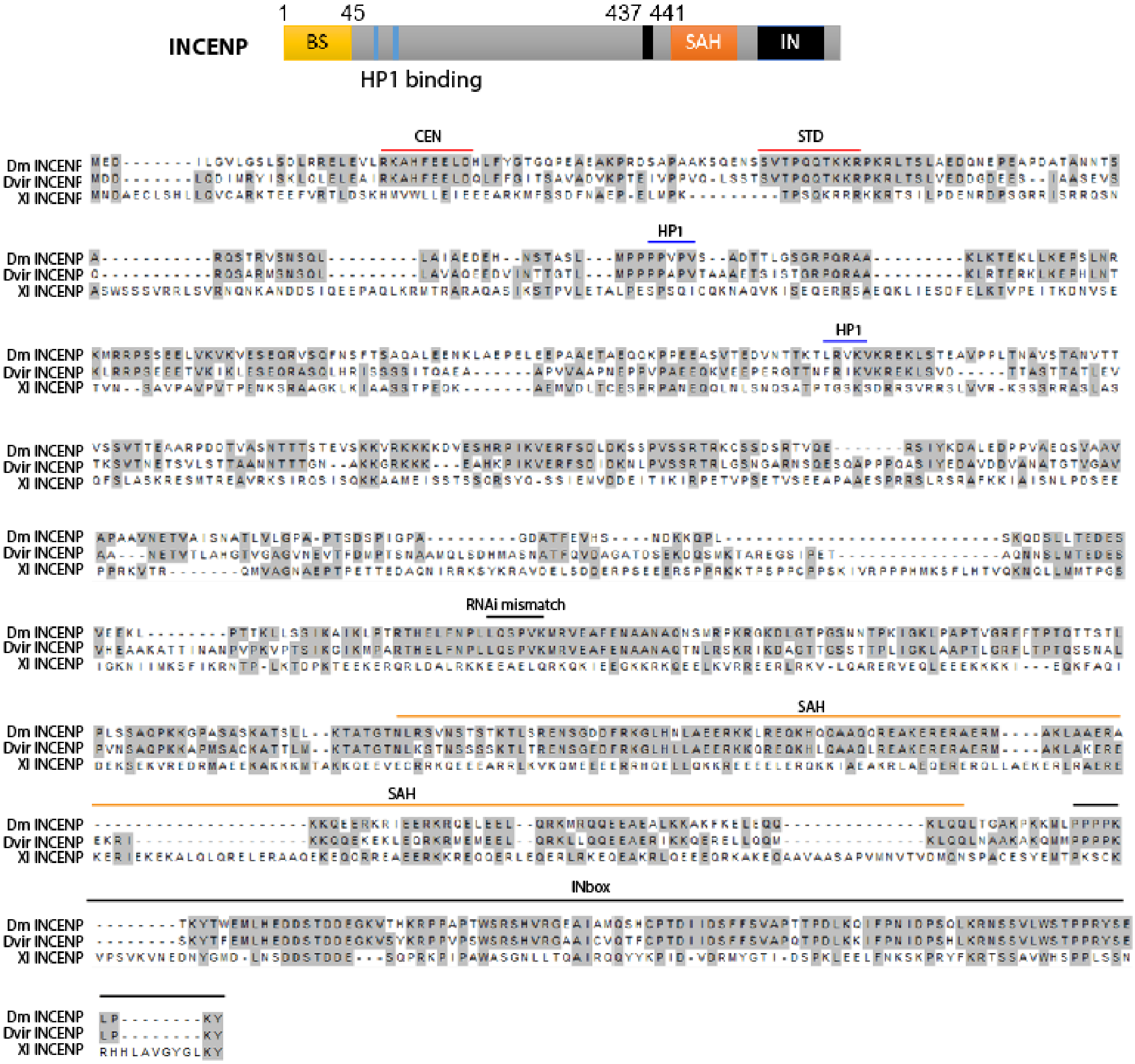
1
45
437 441
INCENP
BS
SAH
IN
HP1 binding
CEN
STD
HP1
HP1
RNAi mismatch
SAH
SAH
INbox
Dm INCENP
Dvir INCENP
Xl INCENP

Figure S2

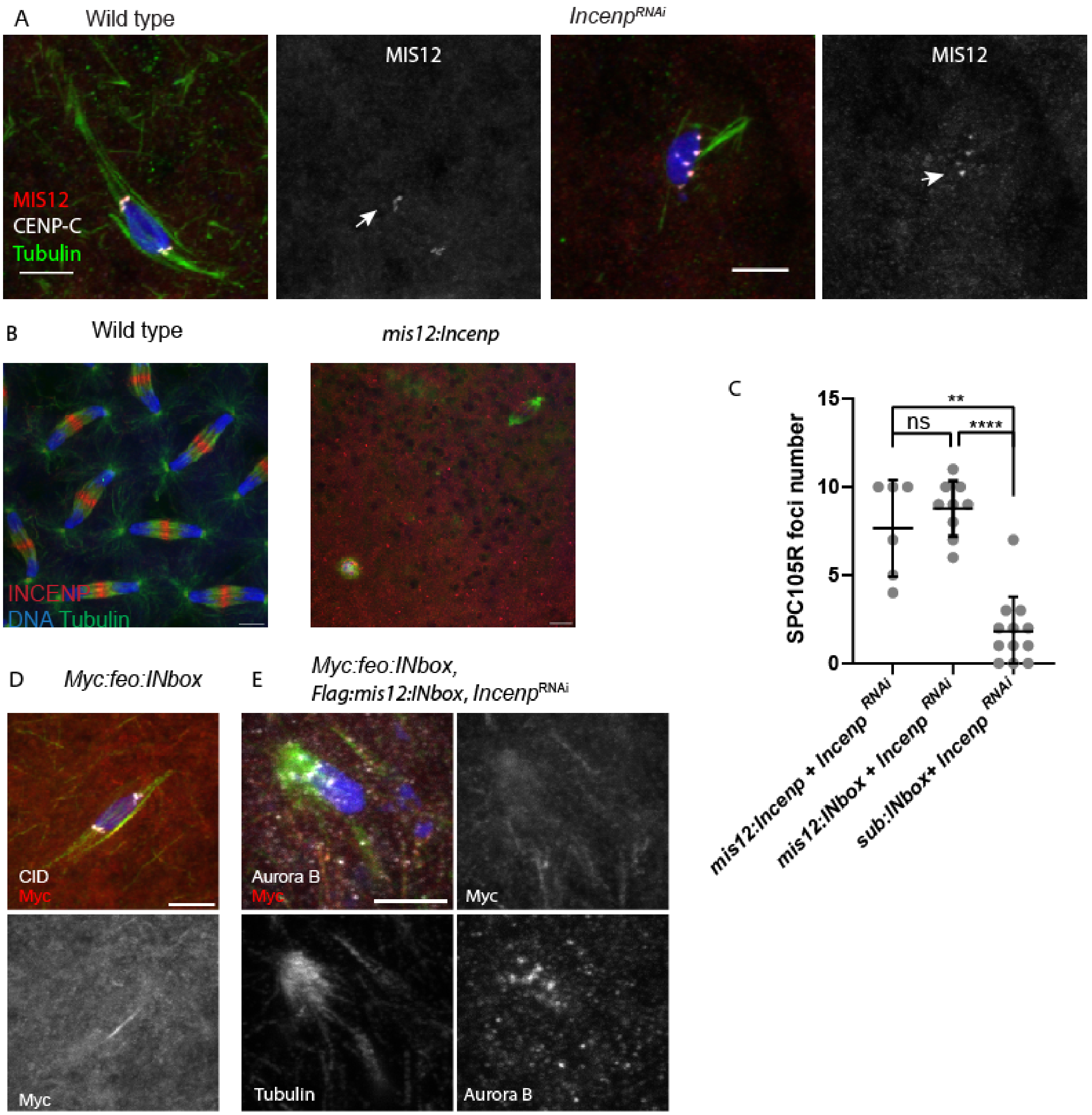
A
Wild type
Incenp^RNAi
MIS12
MIS12
MIS12
CENP-C
Tubulin
B
Wild type
mis12:Incenp
INCENP
DNA Tubulin
C
SPC105R foci number
15
10
5
0
ns
**
****
mis12:Incenp + Incenp RNAi
mis12:INbox + Incenp RNAi
sub:INbox+ Incenp RNAi
D
Myc:feo:INbox
CID
Myc
Myc
E
Myc:feo:INbox,
Flag:mis12:INbox, Incenp^RNAi
Aurora B
Myc
Myc
Tubulin
Aurora B

Figure S3

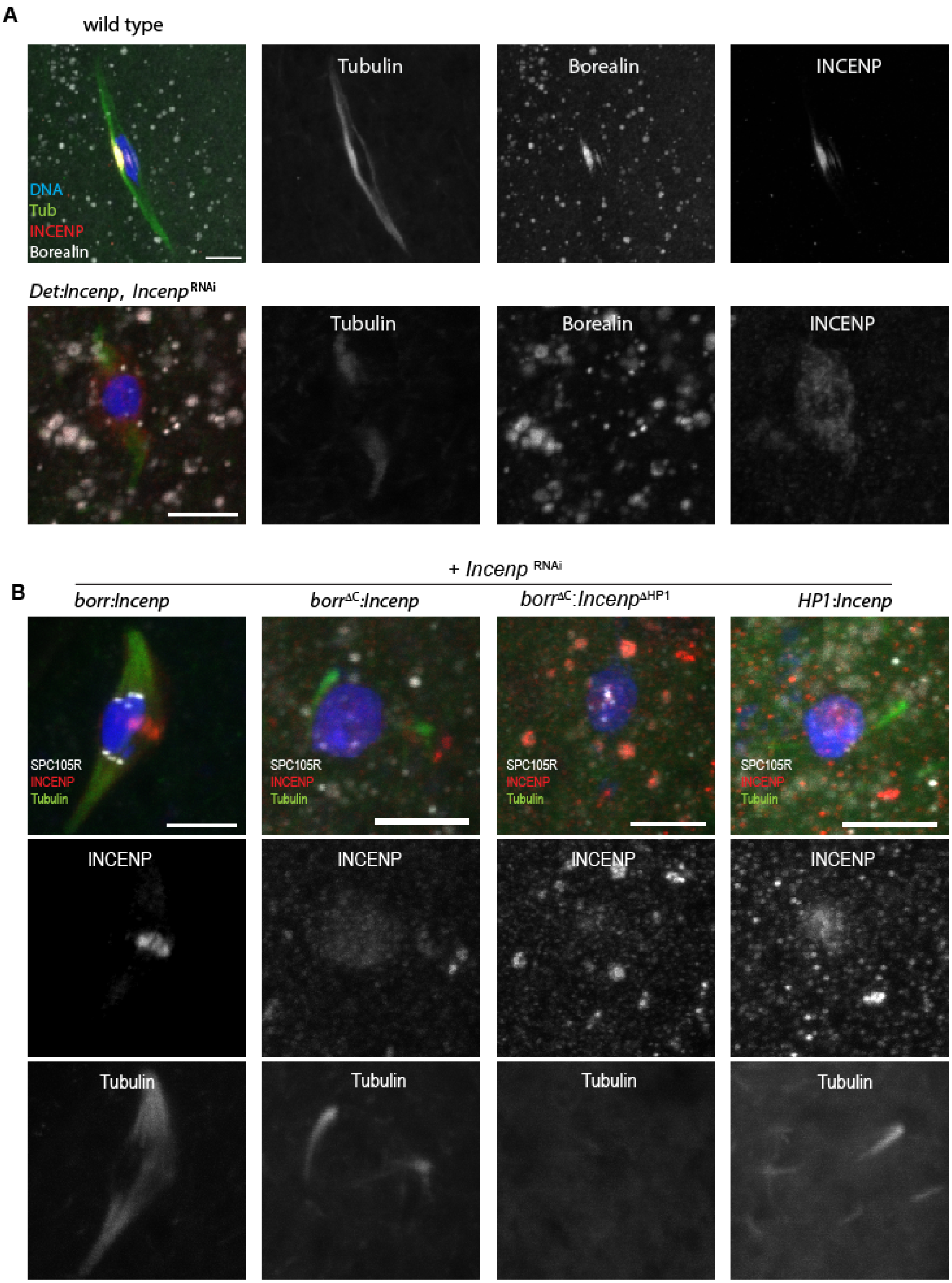
A
wild type
DNA
Tub
INCENP
Borealin
Tubulin
Borealin
INCENP
Det:Incenp, Incenp RNAi
Tubulin
Borealin
INCENP
+ Incenp RNAi
B
borr:Incenp
borrΔC:Incenp
borrΔC:IncenpΔHP1
HP1:Incenp
SPC105R
INCENP
Tubulin
INCENP
Tubulin

Figure S4

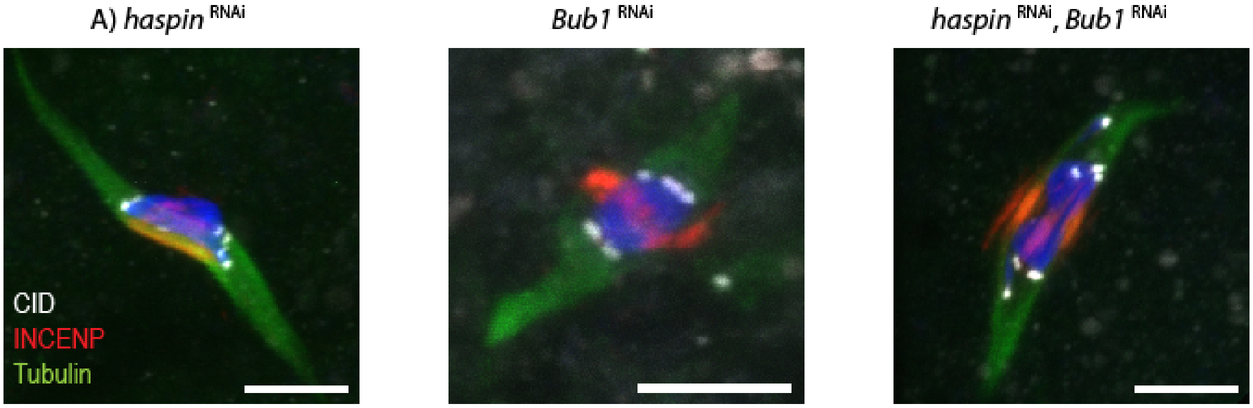


B)

| Genotype | XX | XY | XXY | XO | % NDJ |
|---|---|---|---|---|---|
| ***wild-type*** | 367 | 391 | 1 | 1 | 0.5 |
| ***Haspin RNAi*** | 429 | 352 | 0 | 0 | 0.0 |
| ***Bub1 RNAi*** | 603 | 593 | 0 | 1 | 0.2 |
| ***Haspin Bub1 RNAi*** | 552 | 432 | 2 | 0 | 0.4 |

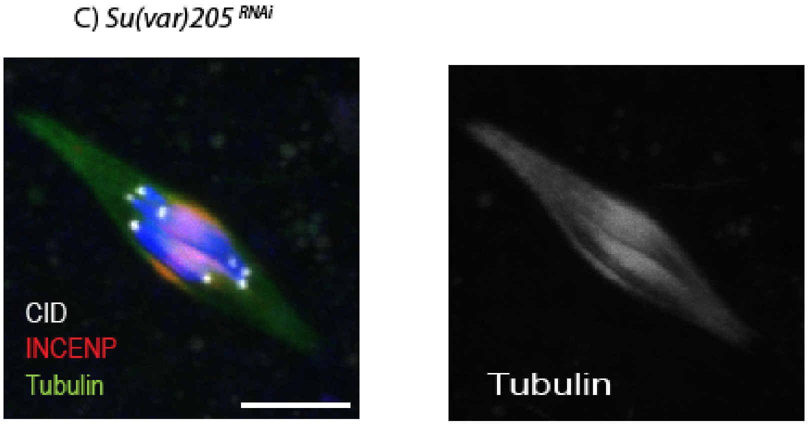
